# Noisy neural systems with static and dynamic Hopf bifurcation parameters

**DOI:** 10.1101/2025.08.04.668381

**Authors:** Priscilla E. Greenwood, Lawrence M. Ward

## Abstract

Stochastic neural oscillations may be quasi-cycles or stochastic limit cycles. Here we discuss how and when these arise in stochastic dynamical systems with Hopf bifurcations, and point to relevant mathematical results. We describe a new type of oscillation, quasi-limit-cycles, which are limit cycles stirred up by noise during time periods called ‘bifurcation delays’ in deterministic models. We define a class of models of neural activity all of which display Hopf bifurcations in their base (fixed bifurcation parameter) deterministic form. We then introduce a two-time, slow-fast, version of our model class, in which the bifurcation parameter, instead of being fixed, changes on a time scale slower than the time scale of the state variables. We demonstrate the effects of the slowly changing bifurcation parameter on the deterministic dynamics of the state variables, in particular ‘delayed bifurcation.’ We find a new understanding about the length of the delay. Most importantly, we display with simulations the effect of noise on the dynamics of both base system and slow-fast system models in our class. Adding noise to a slow-fast model eliminates the bifurcation delay and induces what we call ‘quasi-limit-cycles’ during the delay period. We measure the sizes of the quasi-limit-cycles and show that they are closely related to the sizes of the limit cycles that would arise from the same values of the bifurcation parameter in the deterministic base system. We conclude that, given the similarities of the dynamics of these models under moderate noise, there is little reason to favour one model over another when studying the behaviour of large groups of neurons, i.e., when used as neural mass models.

**Author summary:** Recordings of brain activity display noisy periodic oscillations. Many computational models of this oscillatory activity have what is called a ‘Hopf bifurcation.’ This is a point in the dynamic phase space at which solutions to the deterministic versions of the models change from having a stable fixed point to having a stable limit cycle. In this paper we define a new class of models of oscillatory brain activity all of which have Hopf bifurcations. We compute both deterministic and stochastic path solutions to the models. These solutions give additional insight into an intriguing phenomenon of these models when a parameter slowly changes, called ‘delayed bifurcation,’ as well as expanding our knowledge of their stochastic dynamics. In particular, we define and measure a new type of noisy oscillation, called ‘quasi-limit-cycles,’ that occur during a bifurcation delay in stochastic solutions. The stochastic versions of these models are roughly equally useful as neural mass models.

## 1 Introduction

Most animal brains, including those of humans, display periodic oscillations [1]. There has been some controversy and a lack of understanding about the role(s) such oscillations play, and especially about whether they are causally involved in information transmission in the brain [2–6]. Nonetheless, there continues to be a plethora of theoretical and experimental work investigating their origin and possible functions, e.g., [7–10]. Oscillations, together with fixed points, play an important role in model neural systems, e.g., [11–14]. Many neural and related systems expressing oscillations are well modeled by processes having a Poincaré-Andronov-Hopf bifurcation (henceforth Hopf bifurcation).

A Hopf bifurcation is a critical value of a bifurcation parameter of a two-dimensional dynamical system on one side of which stable fixed point solutions exist for one set of parameter values, whereas for parameter values on the other side stable periodic solutions called limit cycles exist. Hopf bifurcations are relatively well understood in the deterministic regime [15], in the context of comparison at fixed parameter values, although there are still problems worth addressing even there as we will discuss. More recently, however, their dynamic role in particular stochastic systems has been carefully studied, e.g., [16–19], and a more general treatment has appeared [20].

Neuron firing models produce oscillations through dynamic interactions, and many involve Hopf bifurcations. In this paper we describe a class of models of brain oscillations that display a (deterministic) Hopf bifurcation, including a useful novel model, and then characterize the effects of neural noise on the dynamics of these model neural systems. We focus on the behaviour of a class of stochastic systems as a parameter changes, and study the common change in behaviour of the class. This is especially relevant to neural models that have synaptic efficacies as parameters, as these can change over a number of time scales by several different mechanisms [4, 6, 21, 22].

The class of neural models that display Hopf bifurcations that we emphasize contains many of the most popular models of neural activity. This includes the much studied Wilson-Cowan formulation, excitation-inhibition and other networks, and other models with limit cycles. Our prime example is a new, nonlinear, firing model, based on a stochastic linear model of Kang and colleagues [23] that we have studied previously, e.g., [6, 13]. The Kang model is a particularly simple stochastic differential equation system that produces quasi-cycles around a fixed point. As the linear model is unbounded, and thus cannot produce limit cycles, we modify it by imposing the hyperbolic tangent on the active part of the model, much as the logistic transform functions in the Wilson-Cowan model (see later).

In this paper we take a sample path approach to characterizing the behaviour of such neural models, as in [20]. This means that we discuss the dynamics/ stochastic dynamics of the processes themselves, rather than following the Wentzel-Kramers-Brillouin (WKB) approximation approach to exploring these dynamics (cf. [24–26]). Consequently, we are able to introduce a phenomenological way of thinking about the dynamics of the modelled neural systems. Although the stochastic dynamics are our main focus, this needs as a basis an understanding of the underlying deterministic behaviour. The corresponding stochastic analysis is currently unavailable (see the final chapter of [20]), although an analytic approach employing phase-amplitude decoupling via the stochastic averaging method SAM) does address some of these ideas for specific models [19].

Our aim in this paper is to pose, and then, using a path approach, answer a number of questions about a *class* of neural models with Hopf bifurcations, and in particular about their stochastic versions. These questions arise when we consider the role of noise in the brain. Our main points are: (1) We clarify the often-posed question of the relation between stochastic solutions where the model without noise would have a fixed point versus a limit cycle. (2) We investigate the phenomenon of delayed bifurcation in ‘slow-fast’ models with simulations of both deterministic and stochastic paths. (3) We highlight an important role of noise in neural oscillations, in addition to it being necessary for quasi-cycles [13, 27, 28]: not only does it eliminate the delayed bifurcation seen in the deterministic versions of slow-fast models (which was known), but it does this by generating what we call ‘quasi-limit-cycles’ during the delay period in stochastic versions of the models. (4) We measure the sizes of quasi-cycles and quasi-limit-cycles and show how they are related. (5) The outputs of the neural models in our class, which may arise in a variety of ways, are typically indistinguishable in the presence of noise, in a parametric neighbourhood of the Hopf bifurcation.

To our knowledge no detailed comparative study of a class of stochastic neural models that exhibit Hopf bifurcations has appeared, although [19] studies a question similar to ours but about specific examples of what we call “base” models. We define such a class of models of neural activity, which we believe will increase efficiency in addressing what has been an assortment of neural models appearing in various contexts. We describe the context for this class of models, providing some background on dynamical systems that display Hopf bifurcations. This class of neural models is closely related to models of other natural phenomena, such as predator-prey systems [29], chemical reactions such as the Belousov-Zhabotinsky reaction [30, 31] and the Brusselator [32], and indeed to many other systems that are characterized by oscillations. A primary message of this paper will be that when a realistic amount of noise is added to any of these models, the essential differences of their solutions may be reduced or even vanish altogether in a parametric neighbourhood of the Hopf point, so that this aspect of the choice among them is of minor importance. Indeed, to infer from data obtained near the Hopf bifurcation point which member of the model class one is observing will be very difficult if not impossible.

In Section 2 we review the characteristics of deterministic dynamical systems that display supercritical Hopf bifurcations, including the idea of a ‘bifurcation parameter’ that determines whether the system is on one side or the other of a Hopf bifurcation point. In Section 3 we define a class of neural models that display Hopf bifurcations, similar to each other in their structure and basic deterministic dynamics. In addition we develop a simple, representative, new model of neural activity that is a prominent member of this class of dynamical systems. Finally, in this section, we provide simulations of a few of the models in this class that illustrate their similar deterministic dynamics. In Section 4 we exploit the idea of slow-fast, or ‘two-time,’ systems in order to describe the effects of a slowly changing bifurcation parameter on the dynamics of the system. Such a system is said to experience a ‘delayed bifurcation’ relative to the location of the Hopf bifurcation in the set of systems with fixed, distinct, parameter values in established bifurcation theory. Simulations of our new model and some others in the class, together with references to theoretical work, confirm that these models display the delay phenomenon. These simulations also help us to develop a path-related understanding of this phenomenon.

The first five sections of the paper are preparation for Section 5, where we describe how, and cite the mathematics of why, the presence of noise affects the dynamics of the systems in the model class, especially how it reduces or eliminates delay and smooths the transition when a bifurcation parameter crosses the Hopf bifurcation point. We also illustrate some other aspects of the effects of noise in these models, including addressing the origins of the quasi-limit-cycles that occur during delay periods, and presenting results for effects of noise on systems with slowly *decreasing* bifurcation parameters. Finally, in Section 6, we discuss the implications of our results for understanding brain dynamics and other dynamical systems with Hopf bifurcations, as well as directions for future work.

## 2 Hopf bifurcation in 2D deterministic dynamical systems

A zero-centred, two dimensional system of ODEs that exhibits a fixed point at each value of the parameter *a* [33] is:

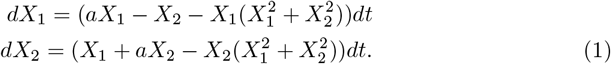

At *a* in a range such that the Jacobian matrix of the system (1) has complex eigenvalues *λ ± iω* the system is said to have a *supercritical* Hopf bifurcation. This occurs in this example at (*X*_1_, *X*_2_) = (0, 0), with *a* = *λ* = 0, *ω* = 1. If *a* = *λ <* 0 the solutions of the state equations display a stable fixed point, and if *a* = *λ >* 0 the fixed point is unstable and the system exhibits periodic oscillations. The system (1) is special in that its ‘bifurcation parameter,’ *a*, is the same as the real part, *λ*, of the eigenvalues, in the linear neighbourhood of the ‘Hopf point,’ *λ* = 0, which separates regions of expansion and contraction. In the class of systems we will consider in what follows, the bifurcation parameter will not be identical to the real part of the eigenvalues of the Jacobian matrix. Rather a chosen parameter of the system, denoted by *S*, will be the bifurcation parameter. It will be responsible for rendering the real part of the eigenvalues of the Jacobian at the Hopf point, *λ*, being either negative or positive, and thus affecting the stability of the system. In some other models, including Hodgkin-Huxley and related neuron firing models, there exist other Hopf bifurcations, called ‘subcritical,’ and showing similar behaviour, that we will deal with later in the paper.

A supercritical Hopf bifurcation of solutions of a system of ODEs occurs because of a difference in solutions between a spiral approach toward a stable fixed point below (*λ <* 0) the bifurcation point (the ‘Hopf point’) and a spiral away from an unstable fixed point and toward a stable limit cycle above (*λ >* 0) the bifurcation point. This difference in solutions depends on the values of one or more parameters of the system. For non-linear systems like (1) the story begins with linearization at a fixed point by computing the Jacobian matrix at that point. The value of the parameter, *a*, at the Hopf point is denoted by *a*_*H*_. In our example, (1), *a*_*H*_ = 0.

The model (1) is useful for illustration because of its symmetry, and because the limit cycles that appear when *a >* 0 are circles with radius 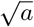. In what follows we will refer to the system (1), and also to the members of the class of related models we specify, as the *base system*, what Su [34] called the ‘frame system,’ and Berglund and Gentz [20] called the ‘frozen’ system. The value of *λ* for any base solution is fixed, and the dynamics will reflect only that one static value. It is important to emphasize this construction in order to avoid confusion between (1) the base system, which comprises a *set* of systems with values of the bifurcation parameter that are different but constant in time for each member of the set, and (2) the associated slow-fast systems to be introduced in Section 4. In the latter, a single system, the bifurcation parameter of the base system becomes a variable, and thus the dynamics is slowly changing according to an additional function of time, from a particular initial behaviour of the system at the initial value of the moving bifurcation parameter.

Figure 1 illustrates the bifurcation dynamics of the base system for (1). The figure shows how the dynamics of the base system are different for values of *λ <* 0 and *λ >* 0. In particular, the figure shows how the solution 𝕏(*t*) = (*X*_1_(*t*), *X*_2_(*t*))^*′*^ of the system *winds in*, as *t* increases with *λ* constant, toward a stable fixed point from arbitrary initial values *X*_1_(0), *X*_2_(0) if *λ <* 0, and *winds out* toward a stable limit cycle from arbitrary initial values *X*_1_(0), *X*_2_(0) inside the limit cycle if *λ >* 0. If the initial values *X*_1_(0), *X*_2_(0) were to be outside the limit cycle the system would wind *in* toward the limit cycle. These interpretations of the dynamics will be of some importance for orientation when we discuss, in Section 4, what is called a ‘slow-fast,’ or ‘two-time,’ system, in which a bifurcation parameter is slowly changed, with a different time parameter, *τ*, so that the related *λ*(*τ*) changes from negative to positive over *τ* time.

**Fig 1.**
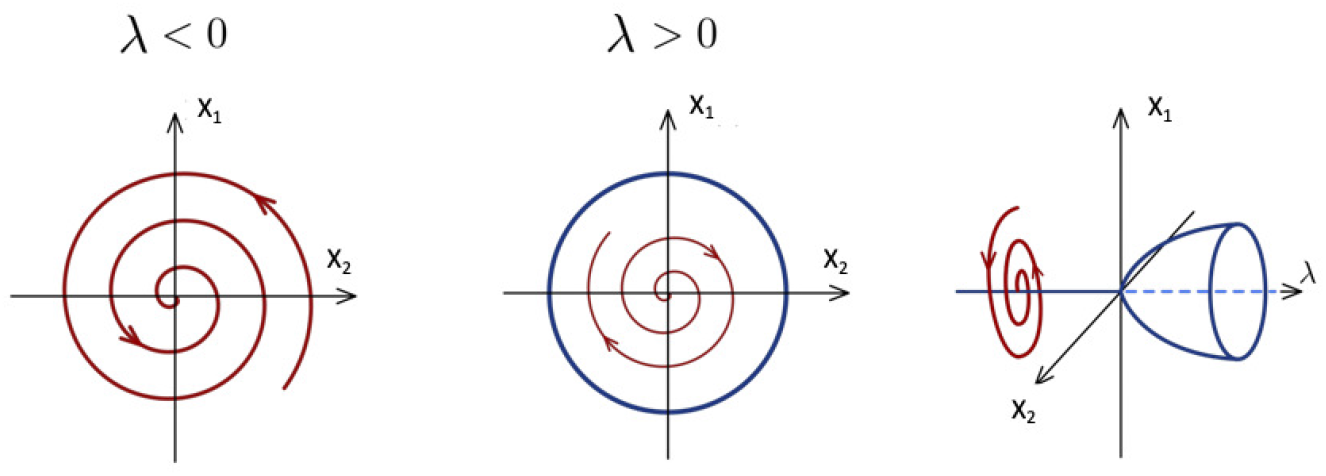
Two dimensional Hopf supercritical bifurcation base system for (1) as a function of *λ*. For *λ <* 0 the red line indicates winding in from an initial position toward a stable fixed point at *X*_1_ = *X*_2_ = 0. For *λ >* 0 the red line indicates winding out from an unstable fixed point, or indeed any other initial values for *X*_1_, *X*_2_. If the model is bounded, as in (1), the winding out is toward a stable limit cycle represented by the blue circle. The rightmost diagram shows how the dynamics differs for values of *λ <* 0 through *λ >* 0. Modified from figure created by Hannes Vogel in Wikipedia article ‘Hopf bifurcation’ under Creative Commons Attribution-Share Alike 4.0 International license.

In the next section we define a class of models in which we are particularly interested because of their common behaviour. Each model in our class will have a supercritical Hopf bifurcation that will be defined in terms of a bifurcation parameter. Their common behaviour, especially that in the presence of noise (see Section 5), will justify using several of them as interchangeable.

## 3 Model class

In this section we define the *base system* of a novel class of dynamical systems that includes a number of more specific models of neural systems. We emphasize the similarities in their dynamics, and use simulations to reveal some more nuanced aspects of that common behaviour. We define the full base system here, including a noise term, but we will develop only the deterministic dynamics in this section, leaving the stochastic dynamics for Section 5. Consider a family of 2 dimensional stochastic differential equations:

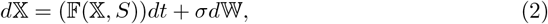

where 𝕏 = (*X*_1_, *X*_2_)^*′*^, 𝔽 = (*f*_1_, *f*_2_)^*′*^ is a nonlinear function that causes the solutions to be bounded; *S* is a bifurcation parameter contained in a bounded interval, *B*, in *R*^1^, and *d*𝕎 = (*dW*_1_, *dW*_2_)^*′*^, a vector of standard Wiener increments with independent components. The scalar *σ* produces the standard deviation of the Wiener increments. 𝔽 not only includes a nonlinearity that bounds the system, but also, critically, it involves positive and negative (or excitatory-inhibitory) interactions between the components *X*_1_, *X*_2_ of 𝕏, which in turn cause the system to exhibit a supercritical Hopf bifurcation. Such an interaction was already apparent in (1).

We define the base systems of this class to be the collection of systems of the form which, with *σ* = 0, are base systems with bifurcation parameter *S* in a particular range, and such that there is a single fixed point that is a Hopf point, *S*_*H*_, of this parameter. In addition the (deterministic) systems are bounded, real, simple, and 𝔽 is differentiable, there is a limit cycle for *S* in *B* ⋂ (*S > S*_*H*_) [35].

For any specific base system of the form (2) with *σ* = 0, having a unique fixed point 𝕏_*S*_, for each *S* in some region of ℝ and a Hopf point at some value *S*_*H*_, we introduce the following notation and specific details:

1. Denote by 𝔸 the Jacobian matrix obtained by linearizing the deterministic part of (2) at a value of *S*.
2. All parameters appearing in 𝔸 except a chosen one, *S*, are fixed. The value of the parameter, *S*, thus determines the eigenvalues of 𝔸, denoted as *λ*_*S*_ *± iω*_*S*_, *ω*_*S*_ *≠* 0.
3. There is a neighbourhood of *S*_*H*_ such that for *S* in this neighbourhood if, *S < S*_*H*_, then *λ*_*S*_ *<* 0, and if *S > S*_*H*_, then *λ*_*S*_ *>* 0.
4. The function *F* in (2) is such that for *S > S*_*H*_, the solution of (2) has a limit cycle |*L*_*S*_| enclosing 𝕏_*S*_. The closer *S* is to *S*_*H*_, the closer *L*_*S*_ is to zero, and *L*_*S*_ converges to 𝕏_*S*_. (| … | denotes ‘size’).

The following are examples of base system models in our model class. In Sections 4 and 5 we will present general results about our model class and use these models as examples. These examples of neural firing models are closely related to neural spiking models, as illustrated by work of Wallace and colleagues [28] for the Wilson-Cowan model, and by several authors for the FitzHugh-Nagumo model.

### 3.1 tanhKang model

A prime example of our model class is what we will call the ‘tanhKang’ model. This model arises from an earlier model developed by Kang et al. [23] and developed further, slightly modified, by Greenwood et al. [13]. The earlier model can be written as

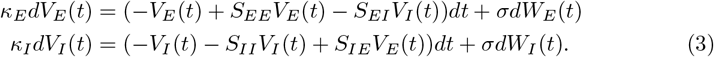

The model describes the time evolution of the local field potentials (LFPs) of excitatory and inhibitory neuron pairs, respectively *V*_*E*_(*t*) and *V*_*I*_ (*t*). The *S*_*ij*_ are synaptic coupling strengths, the *κ*_*i*_ are relative time constants between the excitatory and inhibitory neuron populations, *σ* is noise amplitude, and *dW*_*i*_ are increments of standard Wiener noise processes. If a set of parameters *S*_*ij*_ in (3) yields a Jacobian with complex eigenvalues, *λ ± iω*, as in example (1), the system (3) produces sustained, noisy, oscillations, called quasi-cycles, with frequency distributed in a narrow band around *ω*. The parameters in (3) can be adjusted to give a wide range of noisy oscillation frequencies within the range relevant to brain activity related to mental processes [13].

The model (3) is linear, and, when *σ* = 0, it has a stable solution only for *λ <* 0. For *λ >* 0 the process is unbounded. In order to bound the model for *λ >* 0 we introduce the *tanh* transform as follows. Expressing our new model in the form of (2) we write (3) as

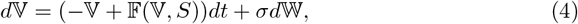

where 𝕍 = (*V*_*E*_*/κ*_*E*_, *V*_*I*_ */κ*_*I*_)^*′*^, 𝔽(𝕍, *S*) = (*f*_*E*_(*V*_*E*_), *f*_*I*_ (*V*_*I*_))^*′*^, *d*𝕎 = (*dW*_*E*_, *dW*_*I*_)^*′*^. Then, as in the model of Wilson and Cowan [36], we have

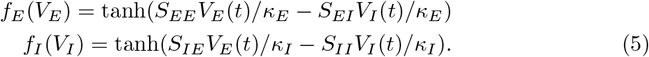

The presence of *tanh* in (5) means there is a limit cycle around *V*_*E*_ = *V*_*I*_ = 0 when *λ >* 0. Note that if we take *η*(*x*) = 1*/*(1 + *e*^*x*^), the standard logistic, and tanh(*x*) = (*e*^*x*^ *− e*^*−x*^)*/*(*e*^*x*^ + *e*^*−x*^), we have tanh(*x*) = 2*η*(2*x*) *−* 1; that is, tanh(*x*) is a rescaled version of the logistic sigmoid function used by [36]. We fix all of the parameters in (4) except for *S*_*EE*_, and choose *S* = *S*_*EE*_ as the bifurcation parameter. The Jacobian for (4) is

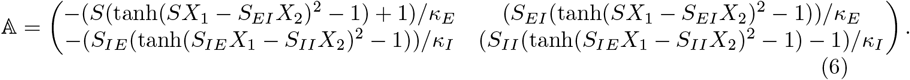

### 3.2 Other members of our model class

Quite a few other models of neural activity belong to our class. They include the much-studied model of Wilson and Cowan [36], which is similar to our new tanhKang model. Here we write the Wilson-Cowan model in a form similar to that expressed by Wallace and colleagues [28]:

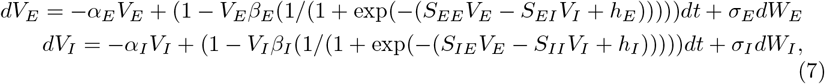

where *V*_*E*_, *V*_*I*_ are excitatory and inhibitory activity, *α*_*E*_, *α*_*I*_, *β*_*E*_, *β*_*I*_, represent rates of transitions from active neuron firing to quiet or from quiet to active firing, *h*_*E*_, *h*_*I*_ are constants, and *S*_*EE*_, *S*_*EI*_, *S*_*IE*_, *S*_*II*_ are synaptic efficacies as in the tanhKang model, comprising the parameters of the model. Wallace et al. [28] showed that various values of *S*_*EE*_, *S*_*EI*_, *S*_*IE*_, *S*_*II*_ yielded solutions with either a fixed point or periodic oscillations, suggesting a Hopf bifurcation. The model becomes stochastic when *σ*_*E*_, *σ*_*I*_ *>* 0. In this form, the model produces quasi-cycles in the fixed point regime, and noisy limit cycles in the periodic regime [19, 28]. Holding all other parameters constant, we can choose *S* = *S*_*EE*_ as the bifurcation parameter and investigate, in terms of the slow-fast model introduced in the next section, how the solutions change as *S*_*EE*_ slowly changes.

Similarly, the Landau-Stuart model with *σ*_*E*_, *σ*_*I*_ = 0 displays a stable fixed point for values of *a <* 0 and stable oscillations for values of bifurcation parameter *a >* 0 [19, 37]:

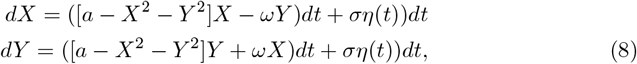

where *X, Y* are excitatory and inhibitory processes, resp., *ω, σ* are constants, and *η* is Gaussian noise. Note that this model is very similar to (1), but with the addition of noise terms. It was used by Dagnino et al. [37] to simulate BOLD activity in the brain in their study of ways to stimulate consciousness in damaged brains.

A model that has been thoroughly analyzed (e.g., [24, 34, 38]) with regard to its supercritical Hopf bifurcation is the FitzHugh-Nagumo model in part of the range of the parameter, *I*. This model is a simpler form of the Morris-Lecar model in part of the range of its parameter *I*, and of the iconic Hodgkin-Huxley model of neural firing in another range of *I*. In the form we use here, it too bears a resemblance to (1), in which a negative cubic term bounds the solutions:

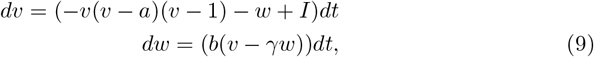

where *v* is voltage across a neuron external membrane, *w* is a ‘recovery current,’ *I* is the applied current, or input, in a particular range, and *a, b*, and *γ* are fixed parameters. In this model, *I* is the bifurcation parameter that determines the value of *λ*, the real part of the eigenvalues of the Jacobian at the Hopf value of *I* = *I*_*H*_, called *I*_*−*_ in [24]. Again, noise can be added to this model, producing quasi-cycles for values of *I* for which *λ <* 0 and noisy limit cycles for values of *I* for which *λ >* 0.

There are forms of the FitzHugh-Nagumo model in another range of the parameter, *I*, and others such as the Hodgkin-Huxley and Morris-Lecar, that contain a *subcritical* Hopf bifurcation, in which the dynamics are quite different from those in the supercritical bifurcation with respect to the eigenvalues of the Jacobian [39]. These models are bistable in an interval of *I* below *I*_*H*_. The behaviour of this subcritical bifurcation version of the FitzHugh-Nagumo model in its quiescent phase, near *I*_*H*_, is essentially the same as that of our model class as studied here. Previous studies of the subcritical Hopf bifurcation of the FitzHugh-Nagumo and related model class are [16, 17, 39]. These models are described in the book of Izhikevich [12].

There are models of other types of systems also related to our class. Prominent among these are predator-prey models such as the Crowley-Martin model [29], which also exhibit Hopf bifurcation points separating parameter regions of fixed points and periodic orbits. Finally, somewhat less relevant to our main points, models of non-living systems, such as the Belusov-Zhabotinsky chemical reaction [30], also display Hopf bifurcations and are related to our class. The B-Z reaction has been modelled by Field and Noyes as a simpler dynamical system [31], termed the ‘Oregonator.’ Several models of bursting neural firing, such as the Morris-Lecar bursting model, are also closely related to our model class.

### 3.3 Deterministic base system behaviour

In this subsection, using simulations, we illustrate the deterministic dynamics of several of the base models in our class for fixed values of *S*. In all of these simulations we employed the Euler method to solve the ODE versions of the equations numerically using double-precision representation in Matlab. This precision allows for 15-16 significant decimal digits and exponents of *±* 308. We ran all of the simulations for 1M time points, with time steps appropriate to the specific model in regard to its stiffness. We should emphasize that our purpose in this study of deterministic models is not to model brain activity, but rather to examine the properties of the several models in order to appreciate the similarities of their stochastic dynamics and slow-fast versions when we come to Section 5.

#### 3.3.1 tanhKang model

Figure 2 displays examples of simulations of the tanhKang base model, (4), with *σ* = 0 and fixed parameter values listed in Table 1. With these parameter values the Hopf point, where *λ* = 0, occurs in this model for *S*_*H*_ = *S*_*EE*_ *≈* 1.5215. The bifurcation parameter *S* = *S*_*EE*_ took the values (A) 1.45, (B) 1.56, (C) 1.6, (D) 1.65, resp., in the four diagrams in the figure. In all cases the initial values for the simulation were (*V*_*E*_, *V*_*I*_) = (0.1, 0.1). At the resolution of the figure, for *S* = 1.45 *< S*_*H*_, *λ ≈ −* 16.67, the winding in toward the fixed point is visible (Figure 2A), but soon disappears into a straight line although mathematically it never arrives at the fixed point. At the end of the 1M time point run the sizes of (*V*_*E*_, *V*_*I*_) are *𝒪*(10^*−*258^). The stable fixed point is never reached via the winding in; there is winding in from any initial value of (*V*_*E*_, *V*_*I*_) other than (0,0), which is degenerate as the system is already at the fixed point in that case.

**Table 1.** Fixed parameters of models used in simulations.

| Variable | Value | Units |
| --- | --- | --- |
| **tanhKang** |  |  |
| $S_{EI}$ | 1.0 | dimensionless |
| $S_{II}$ | 0.1 | dimensionless |
| $S_{IE}$ | 4.0 | dimensionless |
| $\kappa_E$ | 0.003 | seconds |
| $\kappa_I$ | 0.006 | seconds |
| $\Delta t$ | 0.00005 | seconds |
| **Wilson-Cowan** |  |  |
| $S_{EI}$ | 26.3 | dimensionless |
| $S_{II}$ | 1.5 | dimensionless |
| $S_{IE}$ | 32 | dimensionless |
| $\beta_E$ | 1 | dimensionless |
| $\beta_I$ | 2 | dimensionless |
| $\alpha_E$ | 0.1 | dimensionless |
| $\alpha_I$ | 0.2 | dimensionless |
| $h_E$ | -3.8 | dimensionless |
| $h_I$ | -9.2 | dimensionless |
| $\Delta t$ | 0.005 | seconds |
| **FitzHugh-Nagumo** |  |  |
| $a$ | 0.2 | volts |
| $b$ | 0.05 | dimensionless |
| $\gamma$ | 0.4 | dimensionless |
| $\Delta t$ | 0.005 | seconds |

**Fig 2.**
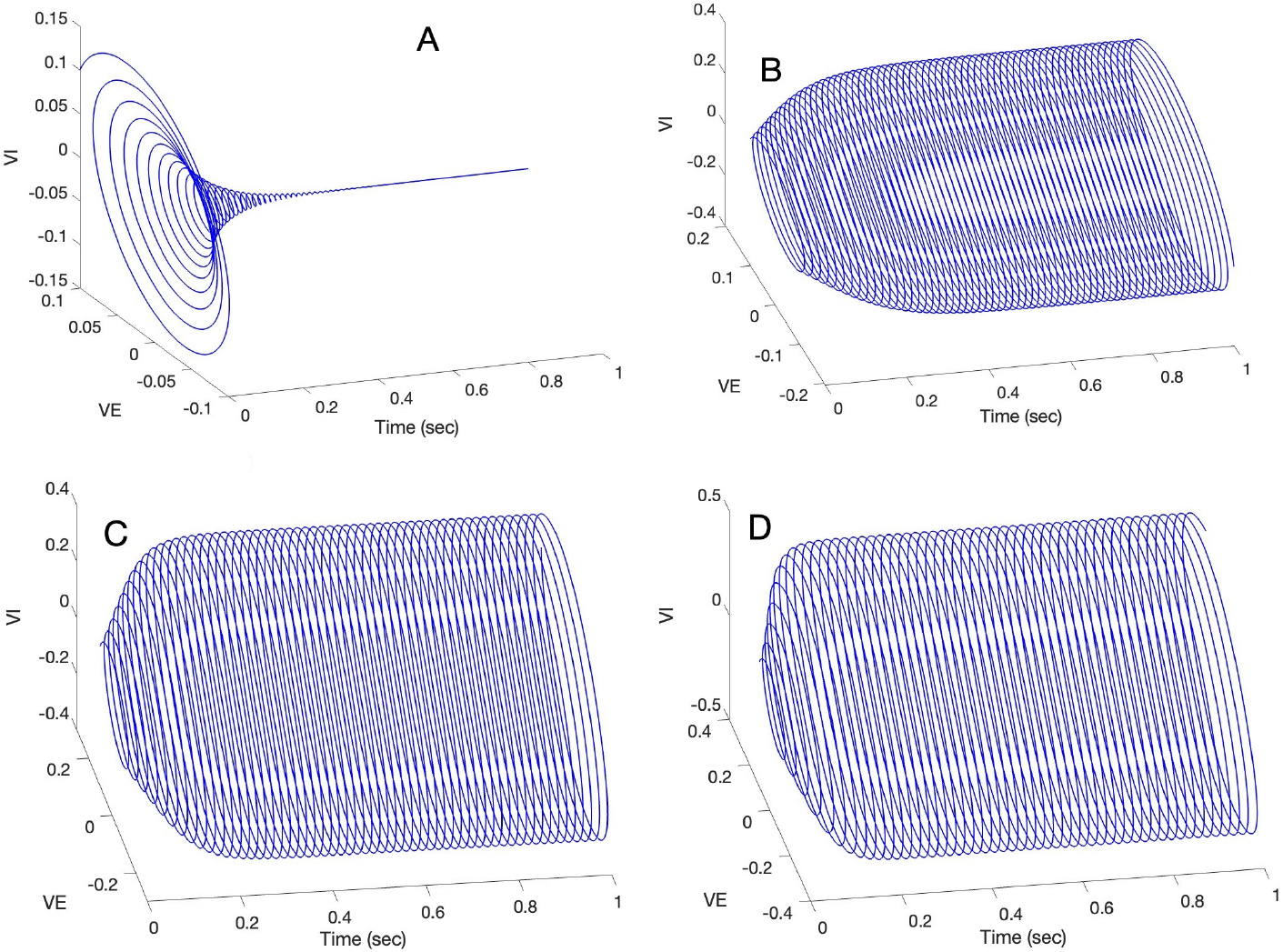
Simulation of tanhKang base model for different bifurcation parameter values and initial values *V*_*E*_ = *V*_*I*_ = 0.1. Only the first 1 sec of the 50 sec run is shown as the remainder of the run is highly similar from 1 sec to 50 sec. (A) With *S* = *S*_*EE*_ = 1.45 the winding in to the fixed point is apparent. (B) With *S* = *S*_*EE*_ = 1.56 the winding out to the limit cycle is apparent. (C) With *S* = *S*_*EE*_ = 1.6 the limit cycle is larger than for *S* = *S*_*EE*_ = 1.56. (D) An even larger limit cycle is seen for *S* = *S*_*EE*_ = 1.65. The limit cycle is larger the larger the distance of *S* from the Hopf point, where *S* = *S*_*H*_ = 1.55. Note also that the larger the value of *S > S*_*H*_, the more rapid is the spiralling out toward the limit cycle.

With *S* = *S*_*EE*_ = 1.56 *> S*_*H*_ *≈* 1.5215, *λ ≈* 1.67 the system displays a limit cycle. With the initial values of *V*_*E*_ = *V*_*I*_ = (0.1, 0.1) the system winds out toward the limit cycle, as illustrated in Figure 2B, although mathematically it never reaches it. The ‘size’ of the limit cycle can be approximated, although it is neither a circle nor even an ellipse, by measuring the maximum chords of the limit cycle for *V*_*E*_ and for *V*_*I*_ resp. once the cycle ‘stabilizes’ to our standards, which means having no noticeable change in size at the scale of the figure. For the limit cycle in Figure 2B the approximate sizes are (*V*_*E*_, *V*_*I*_) = (0.4080, 0.5552). With *S* = *S*_*EE*_ = 1.6 *> S*_*H*_ = 1.5215, *λ ≈* 8.33, the system also displays a limit cycle, as illustrated in Figure 2C. In this case the limit cycle is larger, (*V*_*E*_, *V*_*I*_) = (0.5946, 0.7820), as the value of the bifurcation parameter is further away from the Hopf point. Finally, with *S* = *S*_*EE*_ = 1.65 *> S*_*H*_ = 1.5215, *λ ≈* 16.67, we see an even larger limit cycle, with sizes of (*V*_*E*_, *V*_*I*_) = (0.7746, 0.9830), in Figure 2D. The size of the limit cycle grows as *λ* grows - see property 4 in Section 3. Also, as *λ* controls the rate of approach to the limit cycle, the limit cycle stabilizes sooner for the larger values of *λ*, as can be seen in the figure comparing B, C and D.

#### 3.3.2 WCmodel

Figure 3 displays the results of simulations of the Wilson-Cowan base model, (7), with *σ*_*E*_ = *σ*_*I*_ = 0, and the fixed parameter values listed in Table 1. These values of the bifurcation parameter, *S* = *S*_*EE*_, are near the Hopf point *S*_*H*_ = 24. Figure 3A displays the winding in toward the fixed point, where (*V*_*E*_, *V*_*I*_) *≈* (0.1691, 0.1507), for *S*_*EE*_ = 23 *< S*_*H*_. Figures 3 B, C, and D show winding *in* toward the limit cycle around the unstable fixed point from the initial values of (*V*_*E*_, *V*_*I*_) = (0.1, 0.1), with values of *S*_*EE*_ of 25, 27, and 29, respectively. Similarly to the tanhKang model, these limit cycles are larger for the larger values of *S*_*EE*_: sizes for (*V*_*E*_, *V*_*I*_) are (B) (0.0817, 0.1381); (C) (0.1394, 0.2937); (D) (0.1756, 0.4308). Also the winding in toward the limit cycle is faster the larger is *S*_*EE*_.

**Fig 3.**
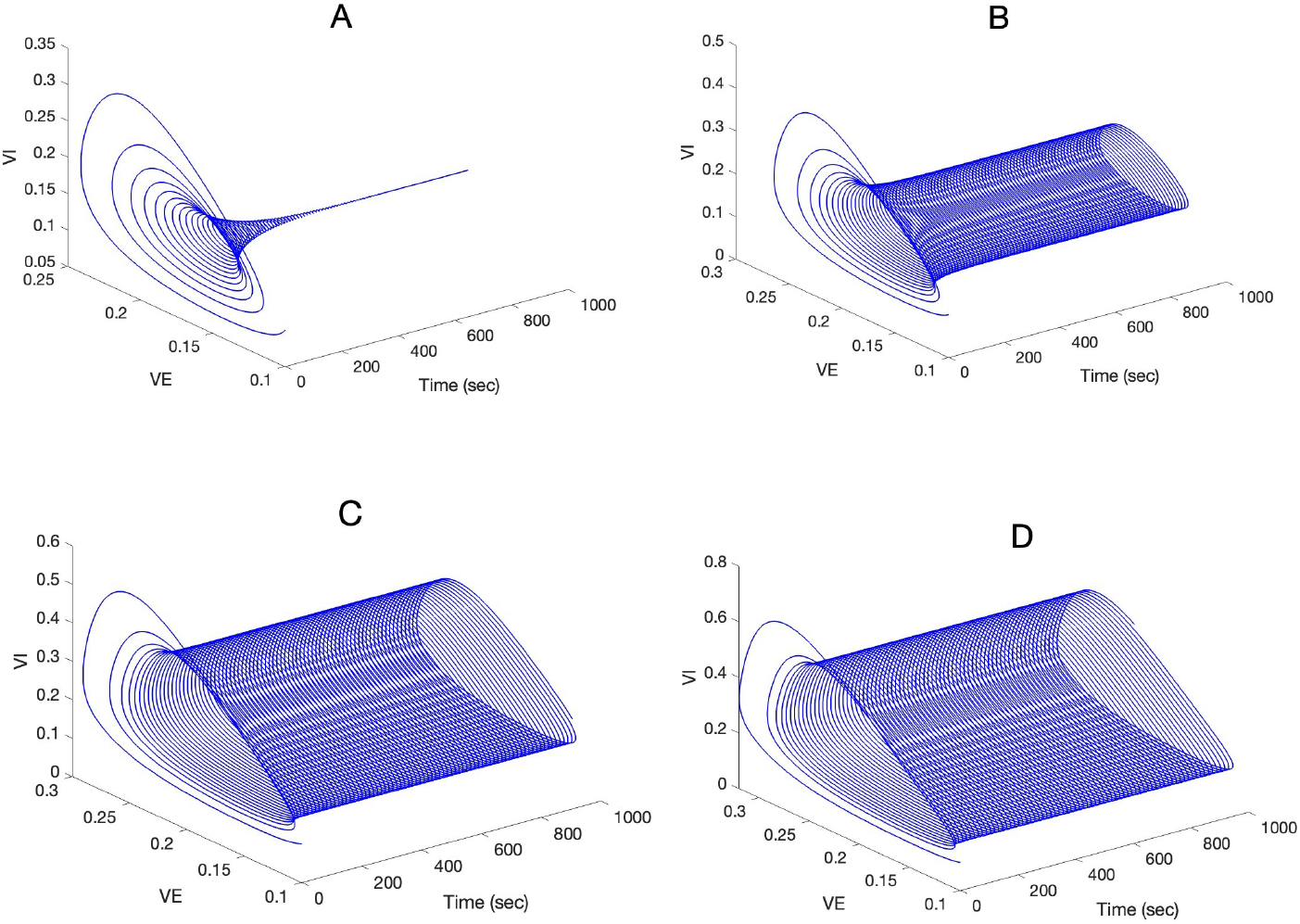
Simulation of Wilson-Cowan base model for different bifurcation parameter values and initial values (*V*_*E*_, *V*_*I*_) = (0.1, 0.1). Only the first 1000 sec of the 5000 sec run is shown as the remainder of the run is highly similar from 1000 sec to 5000 sec. (A) With *S* = *S*_*EE*_ = 23 the winding in to the fixed point is apparent. (B) With *S* = *S*_*EE*_ = 25 the winding out toward the limit cycle is apparent. (C) With *S* = *S*_*EE*_ = 27 the limit cycle is larger than for *S* = *S*_*EE*_ = 25. (D) An even larger limit cycle is seen for *S* = *S*_*EE*_ = 29. The limit cycle is larger the larger the distance of *S* from the Hopf point, where *S* = *S*_*H*_ *≈* 24. Note also the more rapid approach to the stable limit cycle the larger the value of *S > S*_*H*_.

#### 3.3.3 FHNmodel

Figure 4 displays the results of simulations of the FitzHugh-Nagumo base model, (9) with the fixed parameters shown in Table 1. In this model, the bifurcation parameter is *S* = *I*, the current injected into the neuron, and its initial value is (*v, w*) = (0.1, 0.1). With these parameter values, the supercritical Hopf point occurs at approximately *S*_*H*_ = *I* = 0.273 [24]. In this model, the stable and unstable fixed points are increasing functions of *I* [24]. *Figure 4A shows a winding in toward a fixed point, (v, w*) = (0.0964, 0.2410) for *I* = 0.25 *< S*_*H*_ = 0.273. Figure 4B shows a winding in toward a limit cycle for *I* = 0.3, and Figures 4C and D show similar windings in toward limit cycles for values of *I* = 0.4 and *I* = 0.5. The approximate sizes of the limit cycles are (B) (*v, w*) = (0.3937, 0.0934), (C) (*v, w*) = (0.9479, 0.2462), and (D) (*v, w*) = (1.0770, 0.2730) in Figures 4B, C, and D respectively. They are larger for larger values of *I*, as they were in the other models in our class (Property 4 in Section 3). Notice also that the limit cycles are centred around different values of (*v, w*) in Figures 4B, C, and D: (B) (*v, w*) = (0.1235, 0.3127), (C) (*v, w*) = (0.2308, 0.4772), and (D) (*v, w*) = (0.2856, 0.5810). This is because, as mentioned, in this model the coordinates of the unstable fixed points are increasing functions of the bifurcation parameter. This is an unusual view of the FitzHugh-Nagumo model. Some previous studies only considered the periodic behaviour of the *v* variable (e.g., [24, 25, 34]).

**Fig 4.**
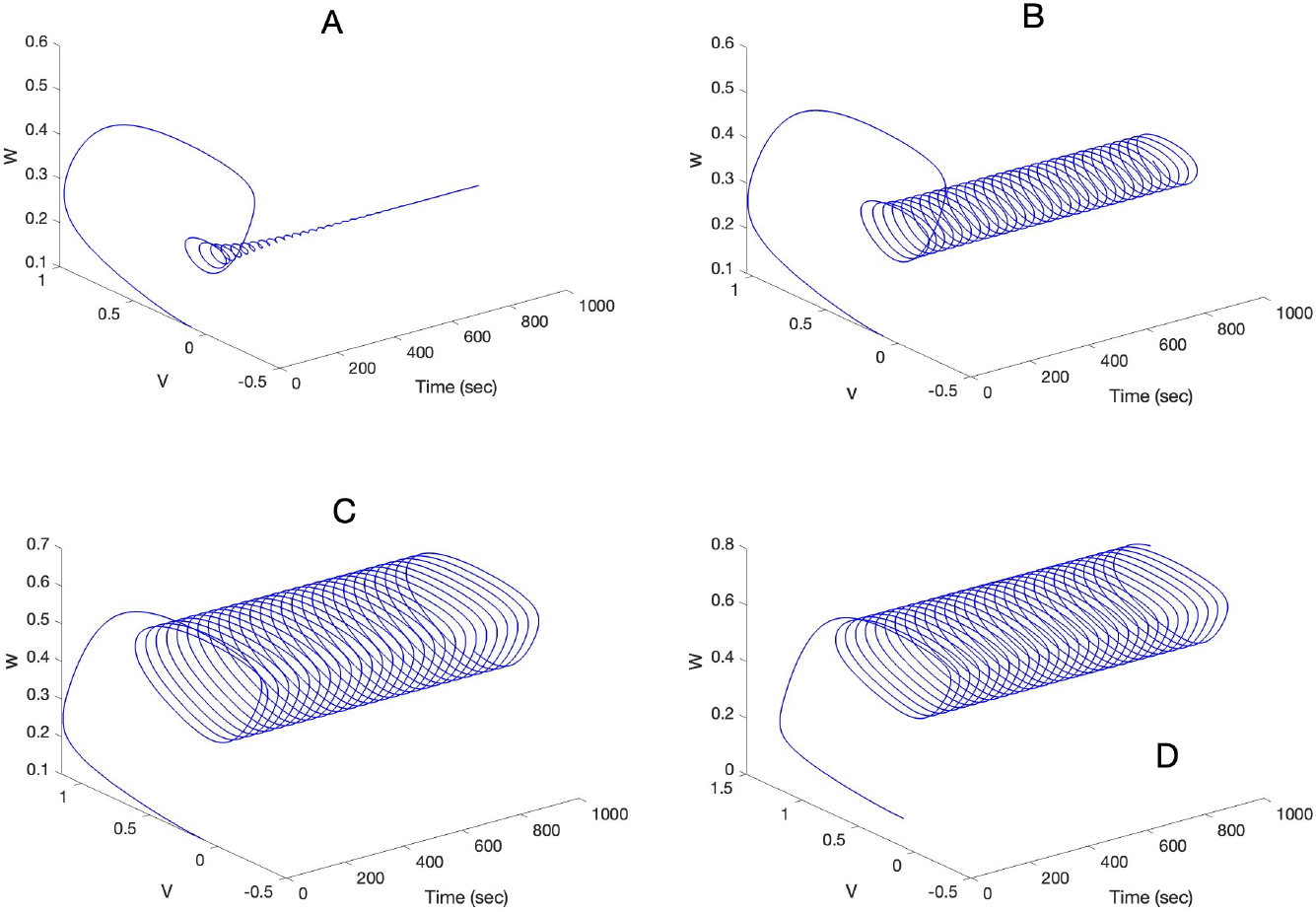
Simulation of FitzHugh-Nagumo base model, (9), for different bifurcation parameter values, *I*, and initial values *v* = *w* = 0.1. Only the first 1000 sec of the 5000 sec run is shown as the remainder of the run is highly similar from 1000 sec to 5000 sec. (A) With *S* = *I* = 0.25 the winding in to the fixed point is apparent. (B) With *S* = *I* = 0.3 the winding in to the limit cycle is apparent. (C) With *S* = *I* = 0.4 the limit cycle is larger than for *S* = *S*_*EE*_ = 0.3. (D) An even larger limit cycle is seen for *S* = *I* = 0.5 (note scale difference). The limit cycle is larger the larger the distance of *S* from the Hopf point, where *S*_*H*_ = *I ≈* 0.273. Note also the more rapid approach to the stable limit cycle the larger the value of *S > S*_*H*_.

## 4 Our class of slow-fast systems: deterministic

A surprising property of dynamical systems that display Hopf bifurcations is the phenomenon of ‘delayed bifurcation’ (e.g., [20, 24, 34]). Because this phenomenon has been observed in natural systems, including neural systems, it is relevant to our model class. In this section we define an extension of our model class to slow-fast, or ‘two-time,’ systems corresponding to our class of base systems (2) again with *σ* = 0, and show with simulations that delay is indeed a property of our slow-fast model class and is not a phenomenon in the base system. We also show that the understanding of delay based on WKB approximations can be enriched by considering high precision simulations of deterministic paths of the state variables. In the following Section 5, we explain the effects of noise on the dynamics of our model class and, in particular, the effect of noise on transitions across Hopf points and on delayed bifurcation. Again, we emphasize that this study of deterministic dynamics is designed to provide a framework within which to develop our subsequent study of stochastic dynamics in these systems, rather than to provide insight into brain dynamics.

### 4.1 Definitions and notation

We define a class of 3 dimensional slow-fast systems in which the first two equations correspond to members of our 2-dimensional class of deterministic (i.e., *σ* = 0) parametric base systems (2). The third equation represents slow change of the value of the parameter *S*, which now becomes the slow variable, *S*(*τ*), in a 3 dimensional slow-fast system, with the process 𝕏 = (*X*_1_, *X*_2_) now becoming the fast process. We adapt the form of [20] Sections (2.2.2)-(2.2.3) to our notation, with their *g* = 1:

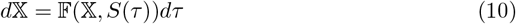

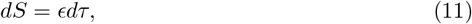

writing the system in fast time with *τ* = *t/ϵ*. It is stated in [20] that one can think of the system (10),(11) as a “perturbation” of the associated base system (2). We emphasize, however, that the slow-fast system, although related to the base system by the first equation, is *not* a version of the base system, but rather a wholly new dynamical system. It is not formed simply by adding the second equation to the base system. The two time scales and the changing bifurcation parameter value are what set it apart from the associated base system.

Because the associated base system has a value of *S* = *S*_*H*_, at which there is a Hopf bifurcation, the question arises: what is the behaviour of the system (10),(11) in an *S*-neighbourhood of *S*_*H*_ ? The answer is in Section 2.2.5 of [20], ‘Hopf bifurcation and bifurcation delay’. The Implicit Function Theorem implies that there is an *S*-neighbourhood of *S*_*H*_ (having the same value *S*_*H*_ in both systems) where 𝔽(𝕏, *S*(*τ*)) = 0, i.e. 𝕏-fixed points of the fast system. We denote by *λ*_*S*(*τ*)_ *± iω*_*S*(*τ*)_ the eigenvalues of the Jacobian matrix *∂*_𝕏_(𝔽(𝕏, *S*(*τ*))), and note that *λ*_*S*_*H* = 0, *λ*_*S*(*τ*)_ *<* 0 if *S*(*τ*) *< S*_*H*_, *λ*_*S*(*τ*)_ *>* 0 if *S*(*τ*) *> S*_*H*_.

Theorem 2.2.12 (‘Dynamic Hopf Bifurcation’) of [20], a version of a theorem of Neishtadt (e.g., [40]), applies to systems in our class (10),(11), telling us that the slow change in the bifurcation parameter, *S*(*τ*), causes *λ*_*S*(*τ*)_ to cross a Hopf point, *S*_*H*_, where *λ*_*S*(*τ*)_ = 0. The change in the value of *λ*_*S*(*τ*)_ from *λ*_*S*(*τ*)_ *<* 0 to *λ*_*S*(*τ*)_ *>* 0 causes the winding in toward the stable fixed point that has been proceeding, ever more slowly as long as *λ*_*S*(*τ*)_ *<* 0, to change to winding out when *λ*_*τ*_ *>* 0. Depending on how long the system has been winding in, with *λ*_*τ*_ *<* 0, producing ever smaller values of 𝕏, and given that the rate of winding out near *λ*_*S*(*τ*)_ = 0 is also slow, it now takes some time before the system actually reaches the region where the attraction to the limit cycle is ‘felt’. The early mathematics about this point is in [34]. Thus, the winding-out solutions of the system stay ‘near’ the fixed point for some time, even while *λ*_*S*(*τ*)_ *>* 0 is increasing, before exponentially ‘jumping’ toward a limit cycle [20]. Figure 5 illustrates this process.

**Fig 5.**
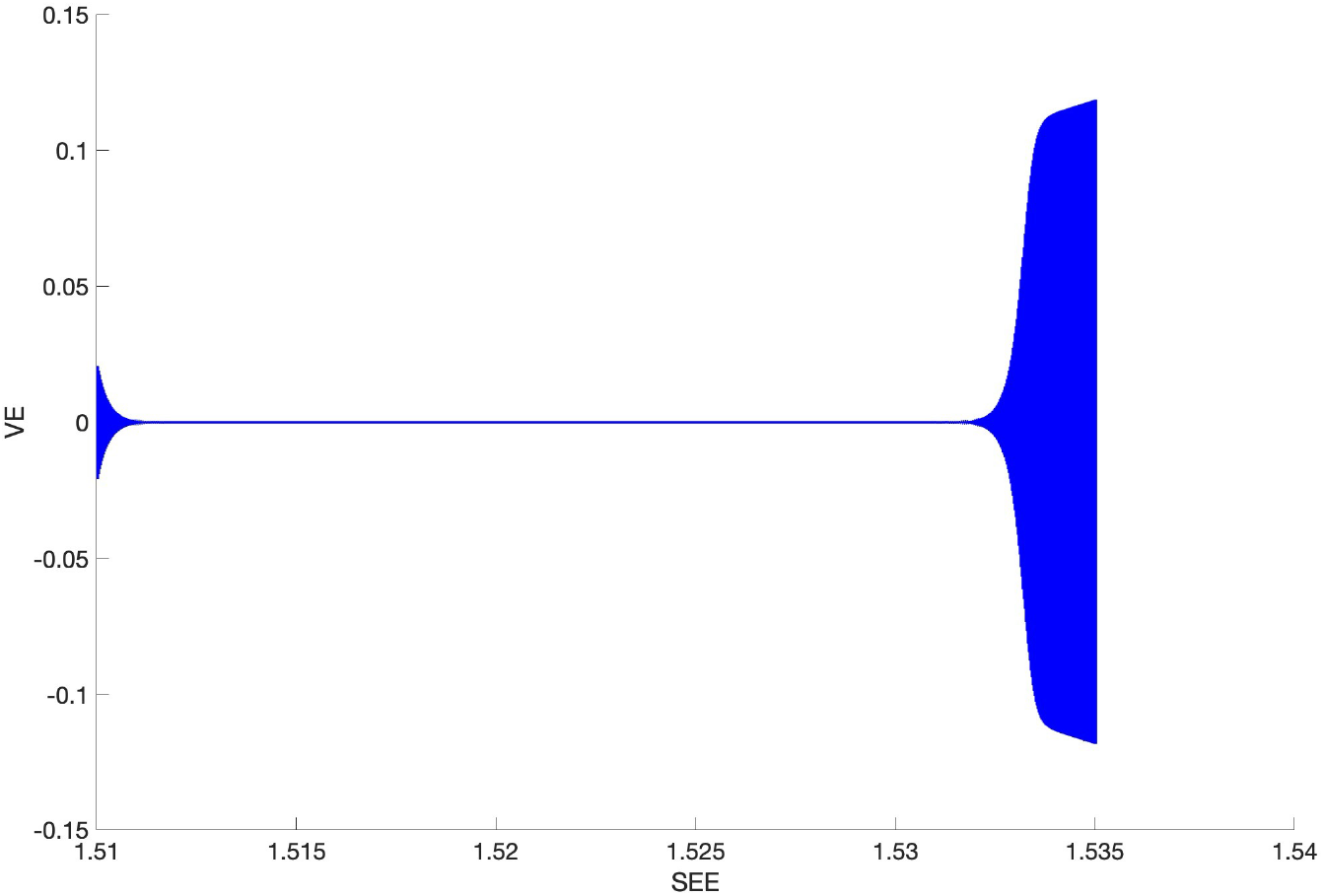
Simulation of a two-dimensional Hopf bifurcation system (tanhKang model) as the critical parameter, *S*(*τ*), is slowly changed, when the initial value of *S*(*τ*) is close to the Hopf point where *λ*_*S*(*τ*)_ = 0. Only one dimension is shown (VE). At left, for low values of the parameter *S*(*τ*), the system winds “down” toward a fixed point from the starting values of *V E* = *V I* = 0.1. Then, at a point somewhat after the point where the value of the parameter *S*(*τ*) causes the real part of the eigenvalues of the Jacobian, *λ*_*S*(*τ*)_, to become positive (*≈* 1.5215, see later), around 1.532, the system executes an exponential escape from the stable equilibrium solution to a stable limit cycle around the unstable fixed point. This behaviour is explained by the analysis and Theorem 2.2.12 in [20].

### 4.2 The delay phenomenon

In our context, as explained above, the slowly changing value of *S*(*τ*) should, according to the base system, cause the system to ‘lose stability’ at the Hopf point of the base system [40], and a stable limit cycle ‘should’ emerge from the resulting unstable fixed point if this were the base system. In the case of a slow movement of the bifurcation parameter, however, in the slow-fast system with *σ* = 0, the solution of the fast system has been observed, and shown analytically (e.g., [20, 24–26]) to stay close to the equilibrium solution of the base system, i.e., to the stable fixed point, until *λ*_*S*(*τ*)_ is considerably larger than 0, before ‘jumping’ toward the stable limit cycle. This phenomenon is termed the *delay of bifurcation*. This has been regarded as a surprising and somewhat exotic result in view of the similarity of the fast system to the associated base system. It is important to emphasize, however, that *this ‘delay’ is relative to the Hopf point computed in the base system*, it is not a ‘delay’ but an inherent property of the slow-fast system. Moreover, the amount of delay (*ceteris paribus*, see later) depends critically on the initial value of the bifurcation parameter, and thus on the initial value of *λ*(*S*(*τ*)) [20, 24, 34, 38]. The initial value of *S*(*τ*), *S*^0^, is such that *λ*_*S*(*τ*)_ *<* 0. Such a value can be determined by adjusting the value of *S* that enters into the Jacobian of the base system until the real part of the eigenvalues of the Jacobian is negative.

Delay of bifurcation, relative to an experiment comprising static values of a bifurcation parameter, has been observed in multiple real systems, including especially squid giant axon [41], thalamic relay nuclei, laser systems, and heating processes, [38], as the bifurcation parameter is slowly changed so that the real part of the complex eigenvalues changes from negative to positive.

#### 4.2.1 History

The now-classical approach to describing bifurcation delay was pioneered by Baer et al. [24] using a WKB approximation to solve the equation system of the FitzHugh-Nagumo model, and to predict both the moment of exponential winding out toward a stable limit cycle and the dependence of this moment on the time since the bifurcation parameter began changing. Later, Su [34] provided a rigorous proof of that approximate result. The approximation stated that when the bifurcation changed slowly and linearly, thus slowly changing the real eigenvalue of the Jacobian from *λ*(*S*) *<* 0 through the Hopf point to *λ*(*S*) *>* 0, the solution did not exhibit large oscillations anytime during the period beginning at *τ*-time zero, at which the 3D process (*X, S*) starts at (*X*(0), *S*(0)), until the first time, *τ*_j_, such that 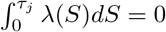. Later, Baer and Gaekel [25] used WKB to obtain approximate solutions for several models for several different ways and speeds at which the bifurcation parameter could change, again emphasizing the integral of the real eigenvalue being equal to zero for large oscillations to occur. Also, Bilinsky and Baer [26] later used WKB to study the effects of a slowly changing bifurcation parameter on spatial delay phenomena in an excitable nerve cable model. In contrast, a path approach is exhibited in Theorem 2.2.12 and its proof in [20], which provides a somewhat different description of the delay phenomenon in the deterministic slow-fast system. It does this in terms of a region around the stable fixed point solution in which the path of the solution resides with a certain probability before it exponentially jumps to large oscillations.

These depictions, however, do not help us to envision the delay in terms of neural processes. They do not address of what the delay actually consists. The mathematical discussions about why the solution resides in the stable region are somewhat subtle, and difficult to relate directly to the changing state variable values. Therefore, we provide here, in an example, a description of the delay phenomenon in terms of the values taken by the dynamical variable 𝕏 = (*X*_1_, *X*_2_), as the value of *λ*_*S*(*τ*)_ changes, recalling that 𝕏 is the voltage of a 2D neural process and *S* is a neuronal input parameter.

#### 4.2.2 Our heuristics

Depending on the value of *λ*_*S*(*τ*)_ in the deterministic (zero noise) tanhKang model, and others with a Hopf bifurcation, the path of (*X*_1_, *X*_2_) either winds in toward a fixed point (*λ*_*S*(*τ*)_ *<* 0) or winds out toward a stable limit cycle (*λ*_*S*(*τ*)_ *>* 0) from wherever it is at a given moment (see Figure 2 for a representation in a base system.) Whenever the value of *λ*_*S*(*τ*)_ in the slow-fast system slowly crosses the Hopf point from negative to positive, the path of (*X*_1_, *X*_2_) doesn’t simply jump toward the limit cycle - *it begins to wind out from wherever it is at that moment*. The nature of these dynamics is such that the path winds in or out around a fixed point (which moves slowly with *λ*) from wherever it starts. It doesn’t simply jump to whatever its characteristic long-term, or base system, behaviour would be. So in the slow-fast model, as the slowly moving bifurcation parameter is changing (and thus changing the value of *λ*_*S*(*τ*)_), and the system crosses the Hopf point, the process is always evolving from some point different from where the deterministic process would end up for that specific value of *S*(*τ*). So it is winding out toward a limit cycle, but it takes time to wind out all the way to the limit cycle, especially because the expansion rate is small for values of 1 *>> λ*_*S*(*τ*)_ *>* 0 near the Hopf point. The slower *S*(*τ*) moves, the slower *λ*_*S*(*τ*)_ changes. Depending on how long the process has been below the Hopf point, which will depend on where it began (how far from the Hopf point), and on how slowly (*τ*) the critical parameter is moving, it will take some (fast) time (*t*) to reach the periodic behaviour seen in the base solutions. This is because the longer *λ*_*S*(*τ*)_ has been below the Hopf point the closer to 𝕏(*S*_*H*_) will be the path of (*X*_1_, *X*_2_) because of the winding in, i.e., the nearer it will be to the fixed point. For example, after 1M time points of the run whose first 1 sec is illustrated in Figure 3A, the value of (*X*_1_, *X*_2_) was *𝒪* (10^*−*258^). In a slow-fast system, the slow rate of expansion after the Hopf point is crossed would require some time for the state variables to reach the level of the limit cycle that is produced in the base solution. This results in a delay in reaching the eventual limit cycle(s), although in fact *‘stability is lost’ the moment that λ*_*S*(*τ*)_ *>* 0, i.e., the winding out commences at that point. At some value of *τ* the effect of 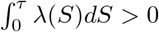 overcomes the accumulated influence of 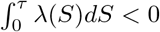 [24, 25, 34]. 𝕏 = (*X*_1_, *X*_2_) has attained a value very close to the fixed point while *λ*_*S*(*τ*)_ *<* 0, and it will take time for the system to climb out of that hole. When this has finished, i.e., when the value of 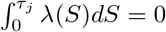, we see the beginning of an exponential winding out, or a ‘jump point,’ toward the limit cycle associated with the current (and increasing) value of *S*(*τ*).

An important point here is that although the integral 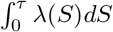 is increasingly negative as the value of *S* increases from *S*(0), and remains negative for a time after *S > S*_*H*_, the winding in or out process of the state variables does not depend on the value of that integral. That process depends on the value of *λ*_*S*(*τ*)_, which will be positive immediately after *S* crosses the Hopf point, *S*_*H*_. The winding out away from the unstable fixed point and toward a stable limit cycle will begin at that time, although that limit cycle will never be reached. Instead, the next change in the bifurcation parameter will cause the process to move toward a different limit cycle, and so on as the parameter changes. This happens regardless of the fact that the integral remains negative for some time after the Hopf point is crossed. Thus, stability is indeed ‘lost’ as *λ*_*S*(*τ*)_ crosses the Hopf point, although that will not be apparent until large oscillations appear. It is, of course, true that the WKB approximation approach [24, 25] identifies the point at which the exponential expansion to the stable limit cycle occurs, as does the path approach of Berglund and Gentz [20]. Moreover, in applications that depend on whether or not large oscillations are present, as in, e.g., lasers, the delay in the appearance of the large oscillations can have practical value. Thus, definitinal stability loss can be dissociated from practical stability loss; both are important but for different reasons.

### 4.3 Examples of the slow-fast model class

#### 4.3.1 Slow-fast tanhKang model

To illustrate the delay phenomenon for the tanhKang model, we simulated solutions to (10), (11) using the tanhKang model with *σ* = 0, as follows (see also Figure 5). We began the simulation with *S*^0^, the critical parameter, at a value that gives *λ*_*S*(*τ*)_ *<* 0, *S*^0^ = 1.4, and moved the parameter value slowly (Δ*S* = 0.00004, *ϵ* = 0.001), solving the tanhKang equations over 1M time points at 0.00005 sec/time point of fast time *t*. In this case, the winding out begins at approximately *S*(*τ*) = *S*_*H*_ = 1.5215 (Figure 6A), i.e., the value of *λ*_*S*(*τ*)_ in the model is close to 0 at this point, and ‘stability is lost’, although this is not visible at the scale of the windings that appear in Figure 6A around *S*_*H*_. In fact, the values of (*X*_1_, *X*_2_) at this time are *𝒪* (10^*−*66^). In this run the exponential ‘jump’ to the limit cycle happened around *S*(*τ*) *≈* 1.644 at the visible scale of the limit cycles (*𝒪* (1)) (Figure 6B), indicating a delay of about 0.1225 from the approximate base system Hopf point to the ‘jump’ point. Looking more closely at the numerics, we see that by *S*(*τ*) = 1.581, *X*_1_, *X*_2_ are at a scale of *𝒪* (10^*−*52^), and at *S*(*τ*) = 1.61 the scale is *𝒪* (10^*−*33^). So the winding out is proceeding, but slowly, toward the ‘jump’ point, but hasn’t gotten there yet. In contrast, when we begin the simulation at *S*^0^ = 1.5, as in Figure 6C, much nearer the Hopf point, the winding out begins at *X*_1_, *X*_2_ values of *𝒪* (10^*−*5^), again at around *S*(*τ*) = 1.5215 as in Figure 6A. It should thus take a much shorter time for the process to attain the values of the base system limit cycles, and indeed that is what happens; the delay in this case is much shorter: from *S*(*τ*) = 1.5215 to *S*(*τ*) *≈* 1.544, or 0.0225.

**Fig 6.**
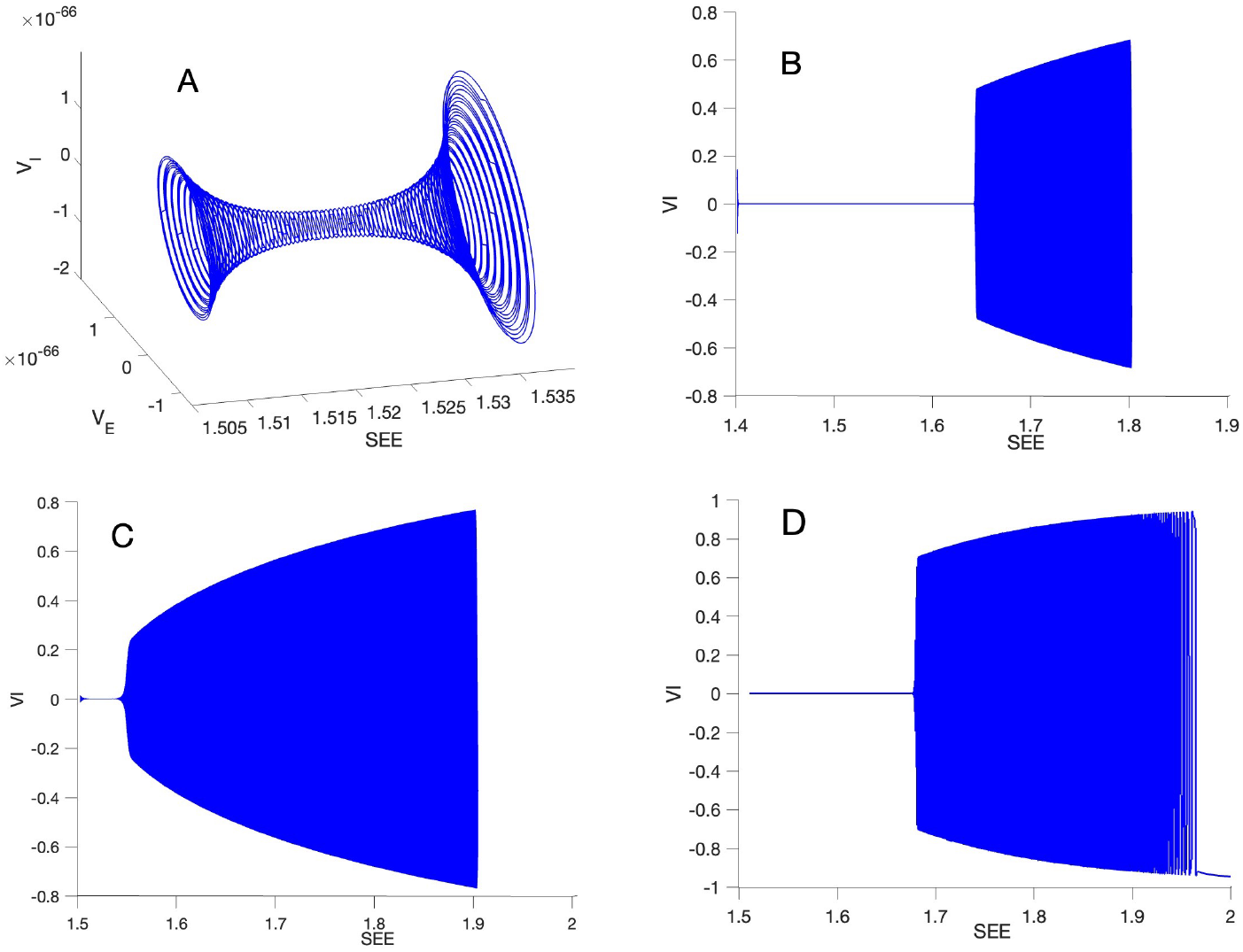
Simulation of (10), (11), using the tanhKang model as the critical parameter, *S*_*EE*_ = *S*(*τ*), is slowly changed across the Hopf point where *λ*_*S*(*τ*)_ = 0. (A) A segment from the simulation in (B) with initial values *V*_*E*_ = *V*_*I*_ = 0.1 and *S*^0^ = 1.4. Here only a small range of values of *S*(*τ*) is shown, bracketing the value of *S*(*τ*) where the winding in changes to winding out, at approximately *S*(*τ*) = 1.5215. Note the scale of the *V*_*I*_ axis: *𝒪* (10^*−*66^). (B) Initial values *V*_*E*_ = *V*_*I*_ = 0.1,*S*^0^ = 1.4. Here a larger range of values of *S*(*τ*) is shown. The ‘jump point’ is approximately 1.644, substantially larger than where the winding out began, displaying a ‘delay’ of approximately 0.1225, based on the single dimension shown (*V*_*I*_). (C) *S*^0^ = 1.5, initial values of *V*_*E*_ = *V*_*I*_ = 0.1. Here *S*^0^ is closer to the Hopf point, which is the same value as in A and B. The ‘delay’ however is much smaller, about 0.0225, with the ‘jump’ point at approximately 1.544. (D) Here the simulation was begun with *V*_*E*_ = *V*_*I*_ = 10^*−*40^, *S*^0^ = 1.51, and the jump point is around 1.68, indicating a ‘delay’ of approximately 0.1585, even larger than when the simulation is begun at *S*^0^ = 1.4 with much larger starting values of *V*_*E*_ = *V*_*I*_ = 0.1.

As predicted by theory [20, 24, 34], the delay depends on the distance in terms of *S*(*τ*), and/or time Δ*τ*, from the base system Hopf point at which the simulation is begun. This result, however, is *ceteris paribus*, that is, all else being equal. Figure 6D displays a result that not only reinforces our analysis in subsection 4.2, but also indicates that the dependence of the delay on the distance of *S*^0^ from the Hopf point is simply a proxy for the values of (*X*_1_, *X*_2_) reached just before that point. In that simulation we began at (*V*_*E*_, *V*_*I*_) = (10^*−*40^, 10^*−*40^), *S*^0^ = 1.51, with Δ*S* = 0.0008. As the Hopf point occurs in this model at around *S*_*EE*_ *≈* 1.5215, the delay should be very small according to the distance dependence alone, similar to that in Figure 6C. In fact, in this simulation delay was quite large, about 0.1585, larger than that in Figure 6B, 0.1225, where *S*^0^ = 1.4 was much further from the Hopf point. What matters for the delay is the value of (𝕏) attained just before the Hopf point is crossed, however those values arise.

#### 4.3.2 Slow-fast Wilson-Cowan model

In simulations of (10), (11) for the Wilson-Cowan model we used Δ*S* = 0.005, *ϵ* = 0.001 and fast time Δ*t* = 0.005. In Figure 7 we illustrate delay in this model. Recall that in the base form of this model *S*_*H*_ = *W*_*EE*_ = 24, so that delay in the ‘jump point’ is measured relative to this value. Figure 7A shows a path with *S*^0^ = 22 displaying a jump point of approximately 25.8 and thus a delay of approximately 1.8 relative to *S*_*H*_ = 24. With *S*^0^ = 23, however, the delay is much smaller (Figure 7B) at about 0.5 because the jump point, although indistinct, is around 24.5. Therefore, the Wilson-Cowan model displays the same behaviour as the tanhKang model, with larger delay the further *S*^0^ is from the base model Hopf point.

**Fig 7.**
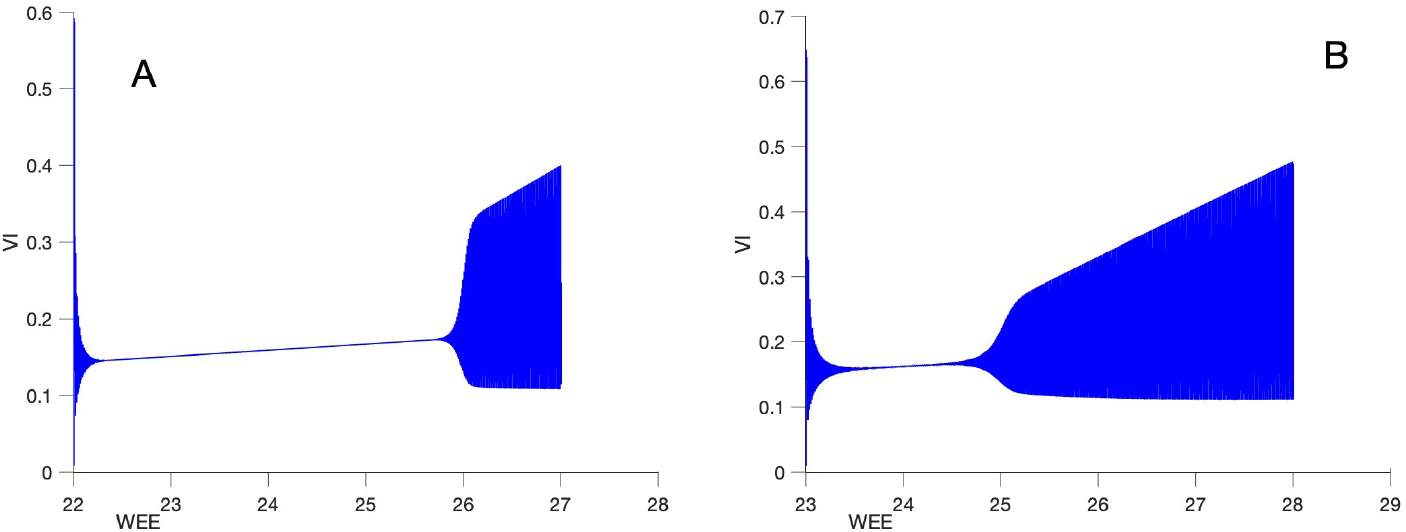
Simulation of (10), (11), using the Wilson-Cowan model, as the critical parameter, *S*(*τ*) = *W*_*EE*_, is slowly changed across the Hopf point where *λ*_*S*(*τ*)_ = 0. (A) Here the ‘jump point’ is about 25.8 as *S*^0^ = 22 is relatively far from the Hopf point at *S*_*H*_ = 24. Thus, the delay is relatively large, about 1.8. (B) *S*^0^ = 23. Here *S*^0^ is closer to the Hopf point. The ‘delay’ is smaller, only about 0.5, with the ‘jump’ point at approximately 24.5.

#### 4.3.3 Slow-fast FitzHugh-Nagumo model

In simulations of (10), (11) for the FitzHugh-Nagumo model we used Δ*S* = 0.0005, *ϵ* = 0.001 and fast time Δ*t* = 0.005. In Figure 8 we illustrate delay in this model. Recall that in the base model *S*_*H*_ = *I* = 0.273, so that delay in the ‘jump point’ is measured relative to this value. Figure 8A shows a path with *S*^0^ = 0.150 displaying a jump point of approximately 0.3815 and thus a delay of approximately 0.2315 relative to *S*_*H*_ = 0.273. With *S*^0^ = 0.22, however, the delay is much smaller (Figure 8B) at about 0.025 because the jump point is around 0.298. Therefore, the FitzHugh-Nagumo model displays the same behaviour as the tanhKang and Wilson-Cowan models, with larger delay the further *S*^0^ is from the base model Hopf point.

**Fig 8.**
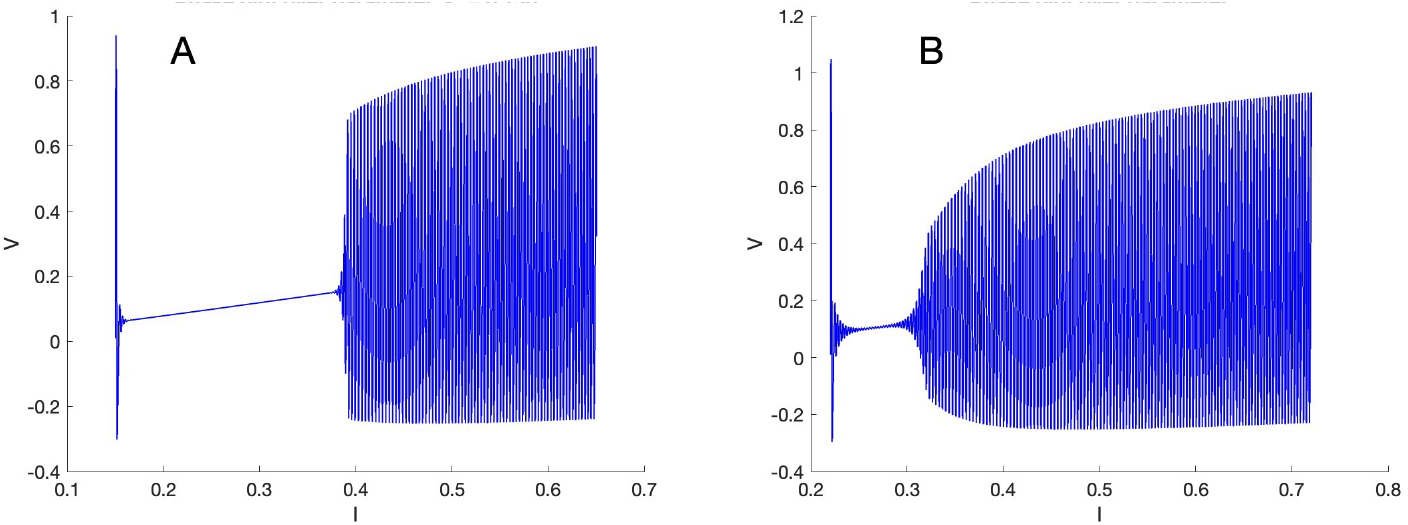
Simulation of (10), (11), using the FitzHugh-Nagumo model, as the critical parameter, *S*(*τ*) = *I*, is slowly changed across the Hopf point where *λ*_*S*(*τ*)_ = 0. (A) Here the ‘jump point’ is about 0.3815 as *S*^0^ = 0.15 is relatively far from the Hopf point at *S*_*H*_ = 0.273. Thus, the delay is relatively large, about 0.2315. (B) *S*^0^ = 0.22. Here *S*^0^ is closer to the Hopf point. The ‘delay’ is smaller, only about 0.025, with the ‘jump’ point at approximately 0.298.

## 5 The effect of noise on static and dynamic Hopf bifurcations

In this section we show by neuron-related examples, and verify by references to general theory, some aspects of the stochastic processes in our model class when noise is included in the model, i.e., *σ >* 0. As we noted earlier, all brain systems are subject to various sources of noise, including noisy synaptic input, channel noise, and so forth from both nearby and more distant areas. Hence the ‘real’ environment of any member of our model class should be taken as ‘noisy.’ For the simulations we use the Euler-Maruyama method for stochastic differential equations. Of particular interest is the understanding of what happens in the static base system models of Section 3 and in the dynamic, slow-fast model class of Section 4.

### 5.1 Results for base systems in our model class

Here we describe simulation results for the base systems of our model class with *σ >* 0, and indicate where proofs can be found in a more general setting. Our noisy base system is

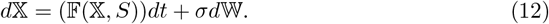

#### 5.1.1 Noisy base system below *S*_*H*_

If *S < S*_*H*_ in a base system in our model class, the solution *X* of (2) has frequency *ω*_*S*_ quasi-cycles centered at 𝕏_*S*_. An example of this for the tanhKang model (4) with *SEE* = 1.45 *< S*_*H*_ *≈* 1.5215 is displayed in Figure 9A, an example for the Wilson-Cowan model with *W*_*EE*_ = 23 *< S*_*H*_ *≈* 24 is displayed in Figure 10A, and an example for the FitzHugh-Nagumo model with *I* = 0.22 *< S*_*H*_ *≈* 0.273 is displayed in Figure 11A. In all cases, the deterministic solution winds in from the initial values of *V*_*E*_ = *V*_*I*_ = 0.1 toward the stable fixed point, whereas the noisy solution produces quasi-cycles around the fixed point. Notice that the noise keeps the quasi-cycles well away from the fixed point. Theoretical results about quasi-cycles can be found in [42, 43]. A theorem in [43] shows that the process represented in Figures 9A, 10A, and 11A are approximated by the product of a rotation and an OU process.

**Fig 9.**
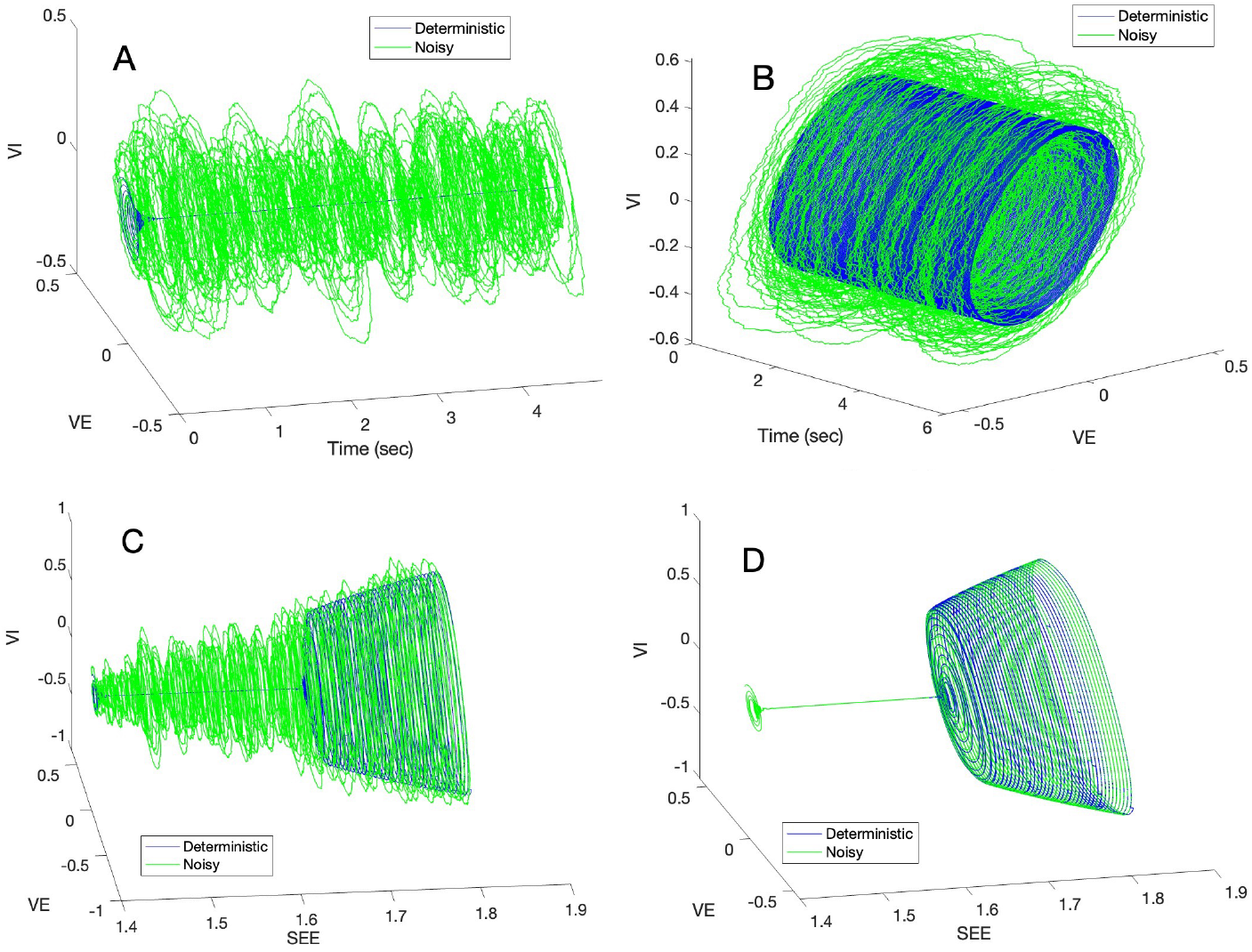
Simulation of tanhKang model (4) with Δ*t* = 0.00005, *S*_*H*_ *≈* 1.5215. (A) Quasicycles with *S*_*EE*_ = 1.45, noise *σ*_*E*_ = *σ*_*I*_ = 0.002. (B) Noisy limit cycles with *S*_*EE*_ = 1.6, noise *σ* = 0.002. (C) Slow-fast system with *S*_*EE*_ = 1.4 to *S*_*EE*_ = 1.8, crossing *S*_*H*_ *≈* 1.5215, Δ*S*_*EE*_ = 0.004, *ϵ* = 0.001, noise *σ* = 0.002. (D) Same as (C) but with noise *σ* = 0.00000000002.

**Fig 10.**
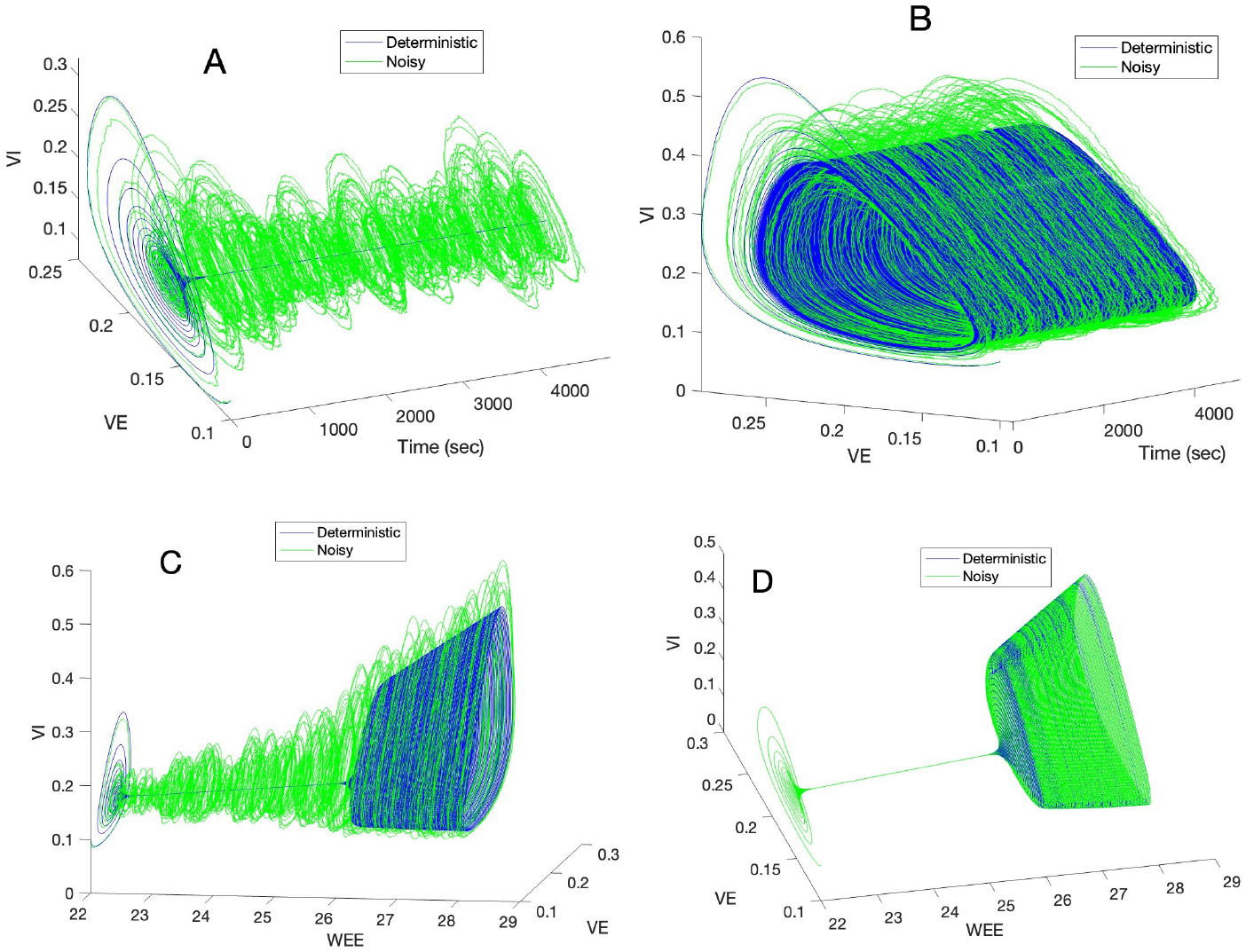
Simulation of Wilson-Cowan model (7) with Δ*t* = 0.005. (A) Quasicycles with *W*_*EE*_ = 23, noise *σ*_*E*_ = *σ*_*I*_ = 0.002. (B) Noisy limit cycles with *W*_*EE*_ = 27, noise *σ*_*E*_ = *σ*_*I*_ = 0.002. (C) Slow-fast system with *W*_*EE*_ = 22 to *W*_*EE*_ = 28, Δ*W*_*EE*_ = 0.006, *ϵ* = 0.001, noise *σ*_*E*_ = *σ*_*I*_ = 0.002. (D) Same as (C) but with noise *σ*_*E*_ = *σ*_*I*_ = 0.00000000002.

**Fig 11.**
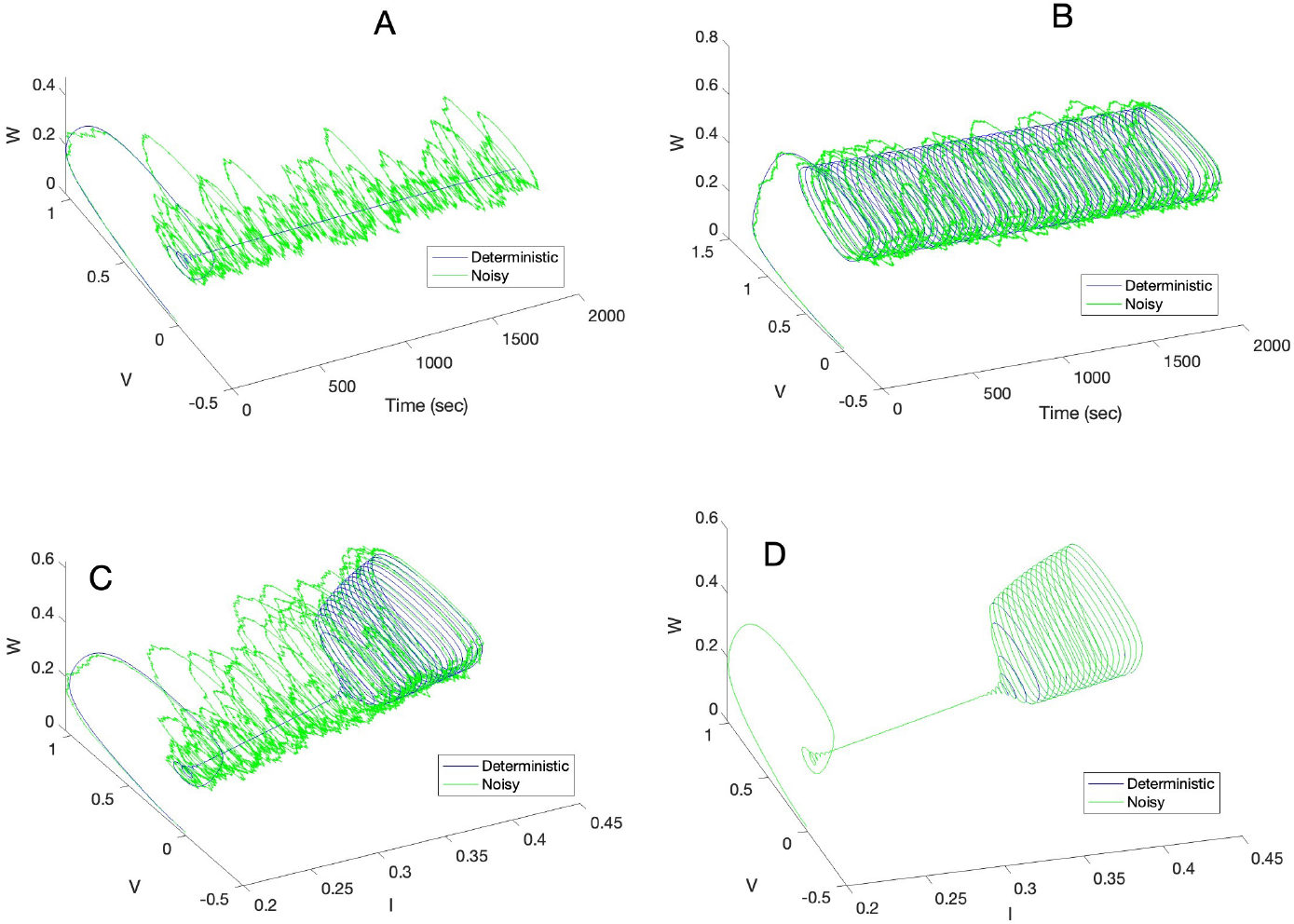
Simulation of FitzHugh-Nagumo model (9) with Δ*t* = 0.002. (A) Quasicycles with *I* = 0.22, noise *σ*_*E*_ = *σ*_*I*_ = 0.001. (B) Noisy limit cycles with *I* = 0.4, noise *σ*_*E*_ = *σ*_*I*_ = 0.001. (C) Slow-fast system with *I* = 0.2 to *I* = 0.4, Δ*I* = 0.0002, *ϵ* = 0.001, noise *σ* = 0.001. (D) Same as (C) but with noise *sigma* = 0.0000000001.

#### 5.1.2 Noisy base system above *S*_*H*_

If *S > S*_*H*_ in a base system in our model class, then the solution, 𝕏, of (2) has noisy limit cycles centred at 𝕏_*S*_. Because the equation for *S* is linear, the sign of *S − S*_*H*_ is the same as that of *λ*, so that we can use the results in, e.g., [20]. An example of this for the tanhKang model (4) with *SEE* = 1.6 *> S*_*H*_ *≈* 1.5215 is displayed in Figure 9B, an example for the Wilson-Cowan model with *W*_*EE*_ = 27 *> S*_*H*_ *≈* 24 is displayed in Figure 10B, and an example for the FitzHugh-Nagumo model with *I* = 0.4 *> S*_*H*_ *≈* 0.273 is displayed in Figure 11B. The deterministic solution winds from the initial values of *V*_*E*_ = *V*_*I*_ = 0.1 toward the stable limit cycle, and the noisy solution produces noisy limit cycles around the unstable fixed point. Notice that with these noise levels the noisy limit cycles stay near to, or actually cycle around, the deterministic limit cycles. Other examples of noisy limit cycles in the Wilson-Cowan model appear in [19, 28]. The dynamics of noisy limit cycles are studied in detail in [44].

### 5.2 Noisy slow-fast system

Here we add *σd*𝕎, *σ >* 0 and not too small, to (10) in the dynamic, slow-fast model of Section 4:

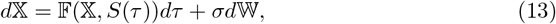

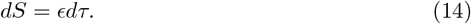

The component 𝕏 = (*X*_1_, *X*_2_) of the solution (𝕏, *S*(*τ*)) has quasi-cycles when *S*(*τ*) *< S*_*H*_ and ‘quasi-limit-cycles’ when *S*_*H*_ *< S*(*τ*) *< S*_*J*_, where *S*_*J*_ (*J* for ‘Jump’) is the value of *S* at which limit cycles become apparent. Note that the ‘jump’ is really an exponential transition toward limit cycles. We use the term ‘quasi-limit-cycle’ because there is no apparent limit cycle present in the deterministic solutions for these values of *S*(*τ*) owing to the delay for *S*_*H*_ *< S < S*_*J*_. In the noisy slow-fast system there is no apparent delay in the transition between quasi-cycles and quasi-limit-cycles, i.e., moderate noise eliminates the delay. For *S > S*_*J*_ limit cycles appear, and the quasi-limit-cycles merge into noisy limit cycles. An example of this for the tanhKang system is displayed in Figure 9C with both the deterministic and a stochastic path included, an example for the Wilson-Cowan model is displayed in Figure 10C, and an example for the FitzHugh-Nagumo model is displayed in Figure 11C. Notice in each figure that the quasi-cycles merge into quasi-limit-cycles and then into noisy limit cycles in the noisy sample path without an obvious transition point, whereas the usual delay is present in the deterministic sample path, with jump points (exponential winding out) in the latter as also shown in Figures 6B, 7B, and 8B. A similar point was made by Powanwe and Longtin [19], although in the context of sets of base models with different (fixed) bifurcation parameter values. They showed that the maxima of stationary distributions of base solutions of stochastic models, such as those we study here (Wilson-Cowan and Stuart-Landau), increase smoothly across the Hopf point for different (increasing) values of the bifurcation parameter.

Theoretical results about the delay times shown in this section, under general conditions satisfied by our model class, can be found both from the WKB approximation [24, 25], and in Section 5.3 of the book of Berglund and Gentz [20]. The results of [20] estimate the delay by bounding the time the solution path stays near the deterministic fixed point. Their results are stated for their more general system 5.3.1 under their assumptions 5.3.1, 5.3.5. The relevant results are stated in their Theorems 5.3.8, 5.3.9, and their following remarks address issues more closely related to our simulation results because of their stochastic path approach. Because of the generality of the setting in [20] it is tedious, and omitted here, to verify that our assumptions in our simpler case do imply their assumptions, so that their results about delay imply our claim that our illustrated results hold for our entire model class. The treatment in [20] includes the reduction and obliteration of delay by noise, but does not address sizes or even the existence of either quasi-cycles or quasi-limit-cycles. However, an asymptotic approximation as *λ →* 0^*−*^ of the size of quasi-cycles can be found in [43].

When there is sufficient noise in both the winding in and the winding out, the noise generates noisy cycles, depending on whether there is winding in around the fixed point as *S* approaches *S*_*H*_ from below, generating quasi-cycles, or winding out toward a limit cycle as *S > S*_*H*_ moves away from *S*_*H*_, generating quasi-limit-cycles. Note that the values computed for *λ*_*S*_(*τ*), the damping or expanding rate (which is determined by *S*(*τ*)), and for *ω*(*τ*), the winding rate, which appear in the eigenvalues of the Jacobian at a fixed point in the deterministic model, (10), (11), also affect the stochastic paths. The sizes of the quasi-limit-cycles are determined by these variables plus the noise amplitude. Just how this happens in the slow-fast system (13),(14), is an open theoretical problem. We believe that something like the result in [43] holds for *λ*_*S*_ *>* 0, and that quasi-limit-cycles are created by the same mechanism as quasi-cycles. In the next subsections we will provide some simulation results that indicate that the members of the slow-fast model class with noise behave similarly for *S*(*τ*) in a neighbourhood of both sides of *S*_*H*_. In addition, there is no visible delay when noise is sufficiently large. The questions of exactly how and why this is the case, what ‘sufficiently large’ noise amounts to, and how the distribution of delay length varies with noise amplitude, apparently have not been sufficiently addressed theoretically (but see [20] for a beginning), and so this is an area that is ripe for future theoretical work.

### 5.3 Noise amplitude

A caveat about the result about noise erasing delay in 5.2 above is that the noise has to be of sufficient amplitude, for it to be true. For very small noise the slow-fast system acts like the deterministic system where *σ* = 0, and so does exhibit the delay phenomenon (cf. [20]). This is illustrated for the tanhKang model in Figure 9D, where the noisy path and the deterministic path are indistinguishable for very small noise *σ* = 0.00000000002. Figures 10D and 11D show the same result for the Wilson-Cowan and FitzHugh-Nagumo models, resp. One nuance of this result is that the rate of change of the slow bifurcation variable, *S*(*τ*), interacts with the noise level to produce it. If the rate of change is very slow, then a very small amount of noise will erase the delay and produce quasi-limit-cycles for values of *S*(*τ*) close to the Hopf point. In Figure 9D the rate of change is relatively fast, Δ*S* = 0.004; slowing it by an order of magnitude, to Δ*S* = 0.0004, results in virtually no delay at all for the same amount of noise (not shown). The distribution of delay length in the stochastic model does not simply scale with noise amplitude when the bifurcation parameter is slowly changing. Note that, in these examples, we have chosen a very small value for *σ*, utilizing the same value for *ϵ* = 0.001 and for the other parameters as in the previous simulations, to illustrate the phenomenon. We have not explored in detail just how largel *σ* must be, for various configurations of parameter values, especially *ϵ* and Δ*S*, to eliminate the delay. The subtle relationship between Δ*S, ϵ, σ* and delay length, other parameters *ceteris paribus*, is a target for further exploration and theoretical development.

Moreover, in our model class, very large noise simply results in random values of 𝕏 regardless of the rate of change of the bifurcation parameter. Thus, there is a version of ‘stochastic resonance’ [45] at play here, with noise in a certain range both erasing delay and obscuring, in the simulations and no doubt also in the neural reality, the transition between quasi-cycles and quasi-limit-cycles. Importantly, the ‘optimum’ range of noise in this sense seems also to depend on the strength of signal being processed by a group of neurons, as well as on the state of arousal in the brain (e.g., [45–47]), and thus could significantly affect the dynamics of these systems. Interestingly, large noise holds surprises for bursting models [48].

We may want to model the “slow” bifurcation parameter, *S*, as being instead a random process, such as an Ornstein-Uhlenbeck process centered at *S*_*H*_, resulting in random alternations between quasi-cycles and noisy limit cycles or quasi-limit-cycles. In this case, the otherwise ‘deterministic’ system (*σ* = 0) would exhibit a variety of behaviours that are a mix of the stable limit point and limit cycle, but only for very short periods of time as the bifurcation parameter, *S*, is constantly changing at random. An otherwise deterministic example for the tanhKang model is shown in Figure 12A, where the value of *S*_*H*_ *≈* 1.5215. In contrast, when *σ >* 0, there are noisy cycles regardless of the value of the randomly changing bifurcation parameter, as in Figure 12B. This possibility has not been explored in theoretical work.

**Fig 12.**
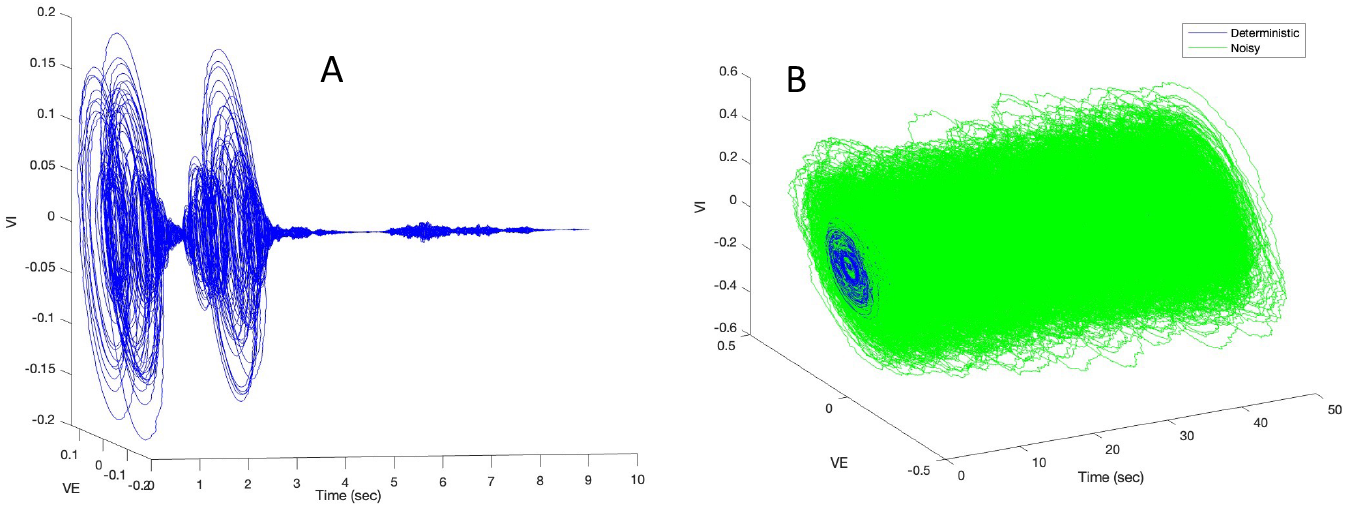
Simulation of tanhKang model (4) with Δ*t* = 0.00005. (A) 𝔹 = 0 but at each time point the value of *S*_*EE*_ changed, *S*_*EE*_(*t*) = *randn*(*t*) + 1.5215, where *randn*(*t*) is a sample from the standard Gaussian N(0,1) and 1.5215 *≈ S*_*H*_ for the tanhKang base system. (B) Same as (A) but with *σ >* 0 as well, *σ* = 0.002.

### 5.4 Noisy cycle size

It is clear from the simulations in Section 5.2 that the sizes of the noisy cycles change as a function of *S*(*τ*). We estimated the noisy cycle size as follows. Because each variable in the two-dimensional fast system oscillates periodically, we estimated the amplitude of the analytic signal, or ‘Gabor amplitude’ from the Hilbert transform [19, 49] on the sample paths of the oscillating variables, computed separately for each component variable, *X*_1_, *X*_2_. In our example displayed in Figure 13A, we simulated the tanhKang model with Δ*t* = 0.00005, Δ*τ* = 0.001, Δ*S*(*τ*) = 0.0008, *S*^0^ = 1.0. To obtain Δ*τ* = 0.001 we kept the value of *S*(*τ*) constant for successive durations of 1000 *t*-time points between increments. We averaged the Gabor amplitude of *V*_*E*_ over each 1000 *t*-time-point episode to obtain a mean Gabor amplitude for each value of *S*(*τ*). These values are the data points displayed in Figure 13B, C and D.

**Fig 13.**
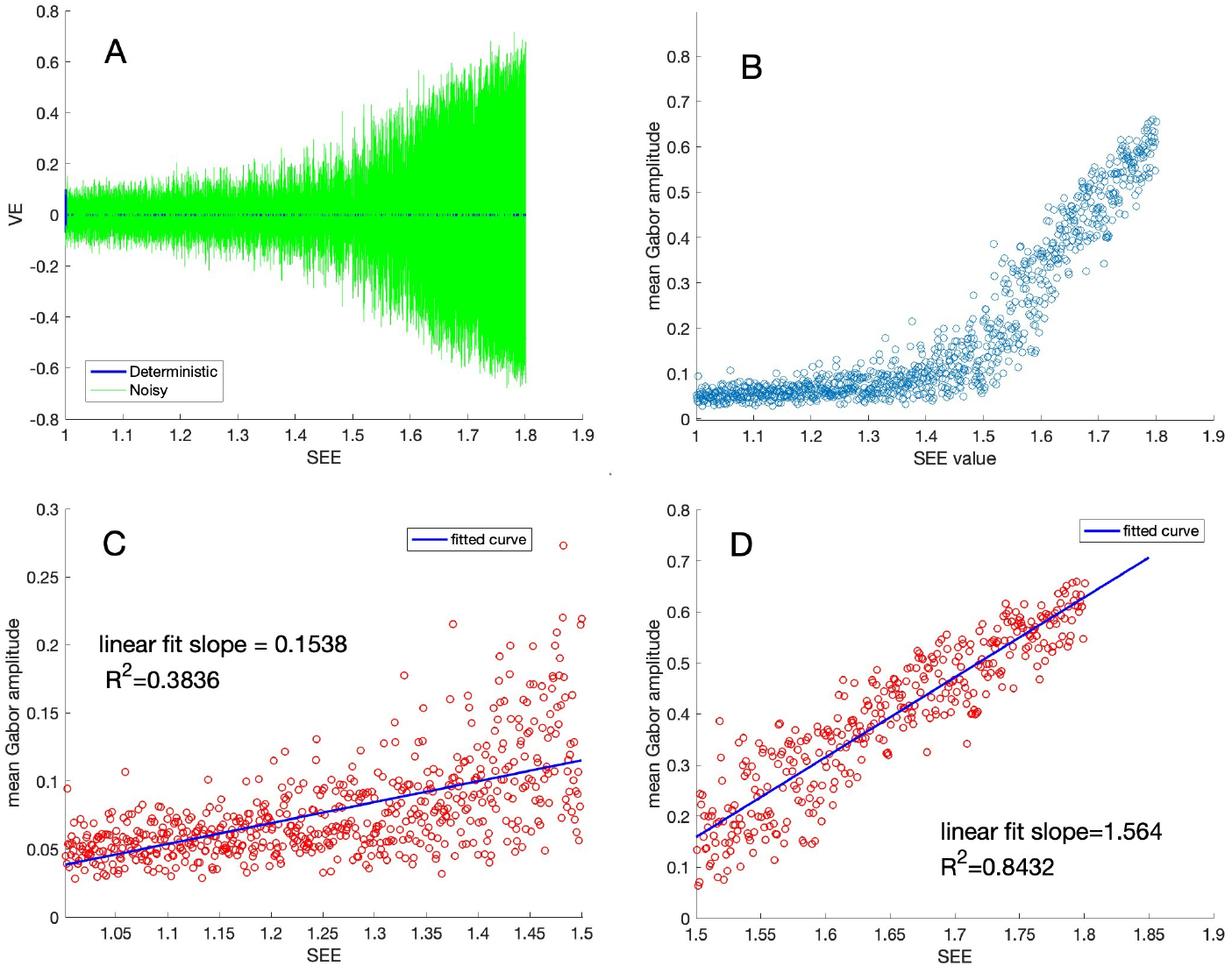
Simulation of noisy (*σ* = 0.002) tanhKang model (4) with Δ*t* = 0.00005, Δ*τ* = 0.001, Δ*S*(*τ*) = 0.0008, *S*^0^ = 1.0. (A) Plot of *V E* sample path versus *S*(*τ*); *V I* is highly similar. Note absence of limit cycles for *S*_*EE*_ *> S*_*H*_ *≈* 1.5215. (B) Plot of mean Gabor amplitudes (see text) versus *S*_*EE*_. (C) Plot of mean Gabor amplitudes for values of *S*(*τ*) *<≈* 1.5, with fit of linear model. (D) Plot of mean Gabor amplitudes for values of *S*(*τ*) *>≈* 1.5, with fit of linear model.

There are several features to note in Figure 13. By choice of initial values (𝕍, *S*^0^), we have arranged that the delay in Figure 13A is so long that large limit cycles do not appear in the deterministic solutions, and we can get a good look at the sizes of the quasi-cycles and the quasi-limit-cycles around the Hopf point, *S*_*H*_, of approximately 1.5215. This is because the simulation was begun at *S*^0^ = *S*_*EE*_ = 1.0, far from the Hopf point (In fact the simulation was started at a point in 3-space, the first two coordinates being the initial values of (*V*_*E*_, *V*_*I*_)). We fitted adjoining lines to two segments of the data, given the obvious elbow and the fact that it occurs at around the Hopf point for this model where quasi-cycles change to quasi-limit-cycles. Figures 13C and D display the resulting two linear fits. The fit for the quasi-cycles, for *S*(*τ*) *< ≈* 1.5 is poor (*R*^2^ = 0.3836), mostly because the wider upward spread of Gabor amplitudes near *S*(*τ*) = 1.5 pulls the fitted line to a larger slope. The slope of the line, 0.1538, is about 1/10 of the slope of the fitted line for the Gabor sizes of the quasi-limit-cycles for *S*(*τ*) *>≈* 1.5, at 1.564. For these values of *S*(*τ*) the fit is much better, *R*^2^ = 0.8432. Notice also that the Gabor amplitudes are particularly variable around the approximate Hopf point. We would expect this from the analysis in [43] as the quantity 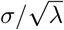 grows as *λ* approaches 0. Indeed, Figure 13B indicates that one might discern an approximate *S*_*H*_ from data via Gabor amplitudes, at least in the case of large delay, whereas doing so from Figure 13A would be more difficult.

Theoretically, it is known that for *S < S*_*H*_ the sizes of the quasi-cycles should grow with 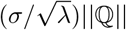 under certain conditions [43]. Here *σ* is the noise standard deviation, ℚ is a normalizing matrix, and *λ ≈* (*S*_*EE*_ *− S*_*H*_) *<* 0. From the way in which quasi-limit-cycles grow in size with increasing values of the bifurcation parameter, however, it would seem that their sizes must be related to the sizes of the deterministic base system limit cycles that would occur for those values of the bifurcation parameter.

We tested this conjecture by computing the relationship between a set of quasi-limit-cycle sizes for the tanhKang model from the scenario in Figure 13, and a set of base system limit cycle sizes with bifurcation parameter fixed at the same values as those of the moving bifurcation parameter, 1.0 *≤ S*_*EE*_ *≤* 1.9. Mindful of the relationship between noise and limit cycle sizes, we did this for several noise levels. The results in Figure 14 A, B show that there is indeed such a relationship, although it depends on the noise level. For large noise (*σ* = 0.01) the sizes of the quasi-limit-cycles are substantially larger than the base system limit cycles, and increase with the value of *S*_*EE*_ at a lower rate. On the other hand, for moderate (*σ* = 0.002) and small (*σ* = 0.0001) noise, the quasi-limit-cycle sizes are virtually identical to the base system limit cycle sizes except near *S*_*EE*_ = 1.5, which is near the Hopf point. Indeed, the stochastic path with small noise in Figure 14 A closely resembles the deterministic path of the tanhKang system when the slow-fast simulation is begun near the Hopf point, as in Figure 6 C. Recall that all of the quasi-limit-cycle results are based on stochastic paths during the delay period because of the very small initial value of the bifurcation parameter.

**Fig 14.**
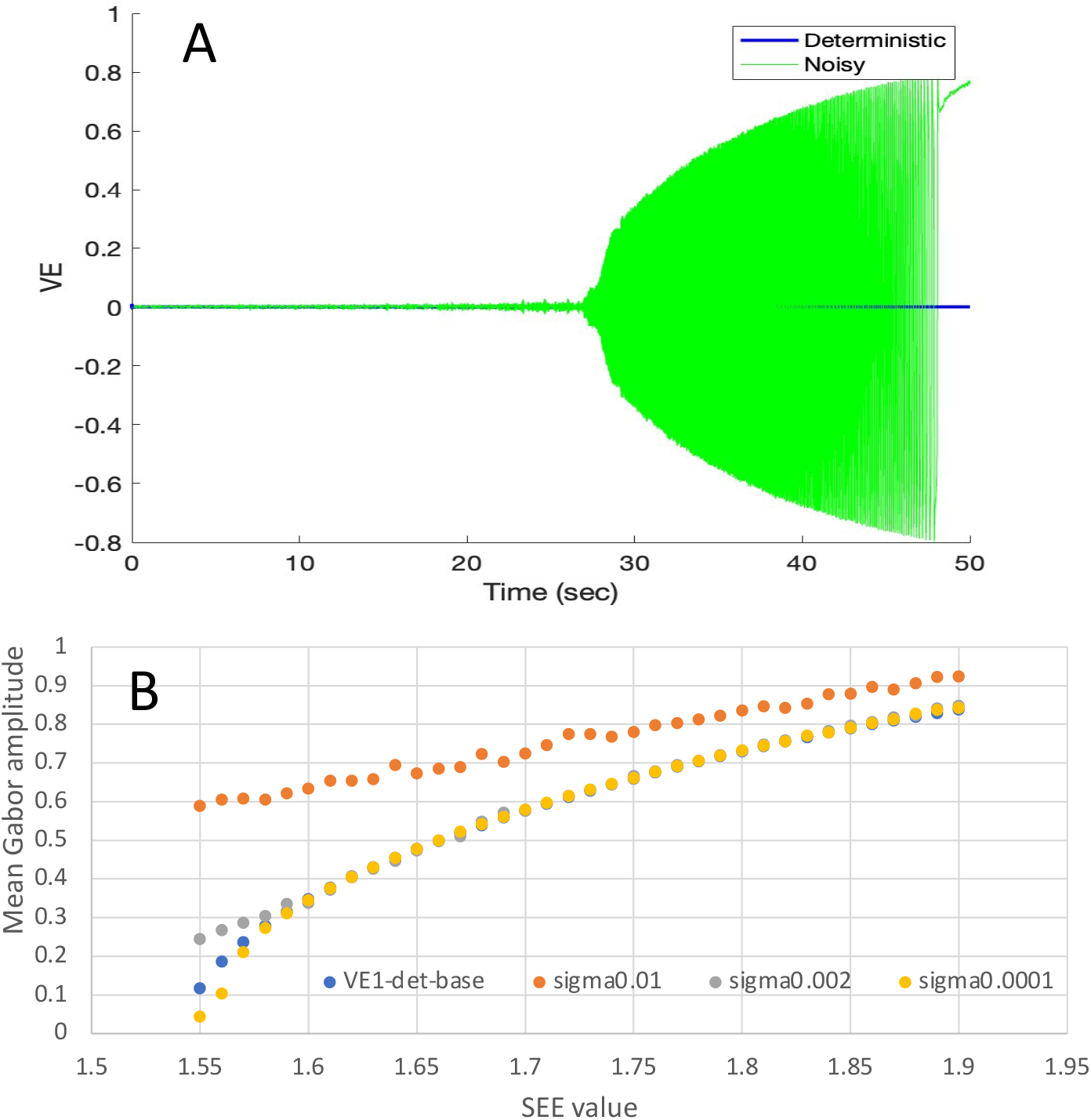
Simulation of noisy (*σ* = 0.002) tanhKang model (4) with Δ*t* = 0.00005, Δ*τ* = 0.001, Δ*S*(*τ*) = 0.0008, *S*^0^ = 1.0, 1.0 *≤ S*_*EE*_ *≤* 1.9. (A) Plot of *V E* sample path with changing bifurcation parameter; *V I* is highly similar. Note absence of limit cycles. In this plot noise *σ* = 0.0001. (B) Plot of mean Gabor amplitudes (see text) of *V E* versus *S*_*EE*_ from 100 instantiations for each of three noise levels and for the deterministic base system. Base system values are the mean Gabor amplitude over 1 M time points for single runs with a fixed bifurcation parameter.

### 5.5 Descending bifurcation parameter

It is understood from various analyses (e.g., [40]) that if the bifurcation parameter in a deterministic dynamical system with a supercritical Hopf bifurcation, such as our class with *σ* = 0, is slowly *descending* in value, causing *λ*(*τ*) to decrease from *λ*(*τ*) *>* 0 above the Hopf point, to *λ*(*τ*) *<* 0 below it, then stable limit cycles persist, while decreasing in amplitude, until the Hopf point is crossed, at which time there is an exponential descent of the persisting limit cycle solutions toward the stable fixed point. There is no delay of bifurcation in this case. However, it is not known what effect noise would have on this process. Figure 15 shows an example for the tanhKang model of a slowly descending bifurcation parameter value both for the deterministic case and for the noisy case with *σ* = 0.002. In the noisy case there is a smooth transition from noisy limit-cycles above the Hopf point to quasi-cycles below it. There is no apparent delay in the transition, and the quasi-cycle and noisy limit-cycle amplitudes are highly similar to both the quasi-limit-cycle and noisy limit-cycle amplitudes for the case where the bifurcation parameter is slowly increasing in value (cf. Figure 13).

**Fig 15.**
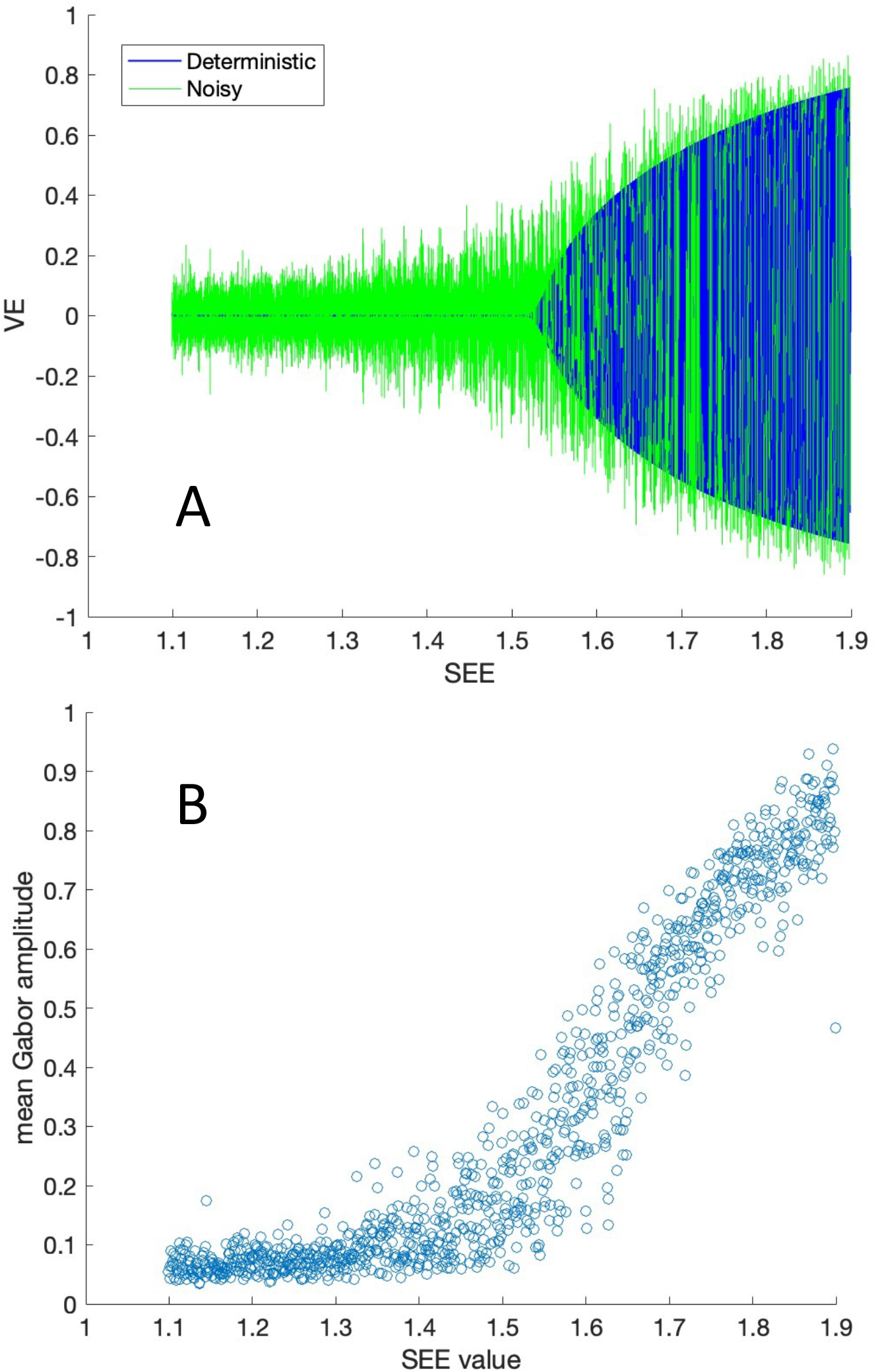
Simulation of tanhKang model (4) with Δ*t* = 0.00005, Δ*τ* = 0.001, Δ*S*(*τ*) = 0.0008, *S*^0^ = 1.9, for *S*_*EE*_ descending in value, 1.9 *≥ S*_*EE*_ *≥* 1.1. (A) Plot of *V E* sample path with changing bifurcation parameter; *V I* is highly similar. In this plot noise *σ* = 0.002. (B) Plot of mean Gabor amplitudes (see text) of *V E* versus *S*_*EE*_. Each point represents mean Gabor amplitudes over 1000 time points at each value of *S*_*EE*_.

## 6 Discussion

Here we summarize the most important takeaways from our work in the form of conclusions, relationships to other work along the same lines, as well as both limitations and future directions.

### 6.1 Conclusions

We have been concerned with characterizing the dynamics of a class of models of neural activity that display Hopf bifurcations. This is because these models generate oscillations, which are arguably important in a variety of neural functions. We began by first reviewing the properties of deterministic dynamical systems with Hopf bifurcations that modulate the transition from stable fixed points into oscillatory regimes. To complete the picture we also described ‘base system’ dynamics, in which values of all parameters of the system are fixed, including that of a bifurcation parameter, and thus the value of *λ*, the real part of the eigenvalues of the Jacobian of the model, and the resulting dynamics, also are uniquely determined.

We explored a class of models of neural activity all of which display Hopf bifurcations in their deterministic form, and thus can generate either winding in toward a fixed point or winding out toward a limit cycle and oscillations. Which dynamics is evident depends on the eigenvalues of the Jacobian matrix, which in turn depend on the values of the model parameters, in particular that of a designated bifurcation parameter. This model class includes several popular models of neural activity, including the Wilson-Cowan model, the FitzHugh-Nagumo model, the Stewart-Landau model, and also a new model, termed the tanhKang model. Using simulations we demonstrated the base system deterministic dynamics for three prominent members of the model class, the tanhKang, Wilson-Cowan, and FitzHugh-Nagumo models.

We then introduced the two-time, slow-fast, version of our model class, in which the bifurcation parameter, instead of being fixed, changes on a time scale slower than that of the other system variables. Using simulations, we demonstrated the effects of the slowly changing bifurcation parameter on the deterministic dynamics, in particular that all of our models displayed delayed bifurcation relative to the Hopf bifurcation point observed in the base system dynamics. We showed that the expected path change from winding in to winding out that should occur at the Hopf point indeed does occur there in the slow-fast system. Moreover, we showed that the delay depends on the values of that state variables that are reached just before the Hopf point is crossed, regardless of how those values are reached, either by a prolonged period of winding in or by adjusting the starting values of the state variables. We also argued, and demonstrated in an example, that approaching the Hopf point from ‘above,’ that is with initial *λ*(*τ*) *>* 0 and decreasing with *τ*-time, did *not* give rise to a delay in the collapse of limit cycles toward the fixed point. We argued that our path approach enriched our understanding of the phenomenon of delayed bifurcation.

We displayed, using simulations, the effect of noise added to the variables on the dynamics of both base system and slow-fast system models in our class. First, adding noise to base system models with *λ <* 0 results in quasi-cycles, whereas adding noise when *λ >* 0 results in noisy limit cycles. Second, adding noise to a slow-fast model during the slow change of the bifurcation parameter across the Hopf point eliminates the apparent delay, as the noisy cycles change from quasi-cycles to what we have named quasi-limit-cycles near the base system Hopf point and then finally to noisy limit cycles at a ‘jump’ point further from the Hopf point.

We showed, using simulations, that the sizes of both quasi-cycles and quasi-limit-cycles (and noisy limit cycles) change with *λ* as |*λ*| moves toward, but do not approach, 0, and appear to be continuous as S moves across the Hopf point. The fact that the slope changes at or near the Hopf point may allow identification of that point in experimental data. We showed that the sizes of quasi-limit-cycles approximate those of deterministic base system limit-cycles for the same bifurcation parameter values until the noise becomes quite large. We also considered the effects of a noisy bifurcation parameter, showing that it caused similar stochastic dynamics even when the remainder of the model was deterministic.

As a general conclusion, we offer the following: given the similarities of the dynamics of these models under moderate noise, which erases bifurcation delay and causes quasi-cycles to merge smoothly with quasi-limit-cycles and the latter with noisy limit cycles, there is little reason to favour one model over another when studying the behaviour of large groups of neurons, i.e., when used as neural mass models. For example, when simulating brain area interactions as done, e.g., by [37], any of the models could be used to similar effect.

### 6.2 Relationship to other work

In this paper, among other things, we have studied and clarified the relationship between (1) the study [43] of quasi-cycles, which are noise-induced cycles from otherwise damped oscillations toward a stable fixed point, and (2) the noisy limit cycles themselves. The quasi-cycles are shown in [43] to be, in distribution, the product of a rotation and an Ornstein-Uhlenbeck process around the fixed point. The noisy limit cycles are shown in [44] to be, in distribution, the product of a winding and an O.U. process around the limit cycle. So we are looking in more detail at the same phenomena and asking: Is there a deeper connection between them? And could this connection lead to a theoretical understanding of the quasi-limit-cycles? One direction that could be taken is that of Powanwe and Longtin [19], who used the stochastic averaging method to decouple amplitude and phase of solutions to the Wilson-Cowan and Stuart-Landau models. This decoupling is similar to that of [43] and [44], and allows analytical results about the relationship between quasi-cycles and noisy limit cycles.

We discovered a natural moving parametric link between fixed points and limit cycles in members of a parametric class of models with Hopf bifurcations, and found ourselves studying slow-fast dynamic models. A neural motivation for the slow parametric link was the slow current in neural bursting models, which are cyclic slow-fast systems studied, e.g., in [48].

Other works studying the role of noise in destroying delays in Hopf bifurcations are [17] for FitzHugh-Nagumo bursters, and [16] for more general bursters. Both [17] and [16] are about subcritical Hopf bifurcations where there is a similar phenomenon and theory of bifurcation delay to that in the supercritical case studied here.

The delay phenomenon has been studied, in both deterministic and stochastic settings, by a variety of authors using various approaches, such as the WKB approximation (e.g., [24, 25]) and a path approach (e.g., [20, 40] on slow-fast dynamical systems. Both approaches address some of the results illustrated here for certain neural models. In particular, the early motivation of the work of Berglund and Gentz was about certain questions in stochastic neural path dynamics, similar to ours. One avenue of further work on our model class would be to continue in the vein of Chapter 5 of [20] to study properties of quasi-cycles and quasi-limit-cycles appearing near Hopf points. A different approach would be to extend stochastic approximation theory arguments like those in [43, 44], but now for our slow-fast stochastic model class (13),(14). Finally, the approach of Powanwe and Longtin [19] could be extended to our class of slow-fast systems. A gap in our theoretical understanding is why the sizes of the quasi-limit-cycles, under small to moderate noise, that appear in the delay time interval between the Hopf point and the appearance of deterministic limit cycles, approximate those of the deterministic base system limit-cycles for the same bifurcation parameter values.

### 6.3 Limitations and future directions

We explored the effects of noise on the dynamics of our model class using simulations. We have found in the mathematical literature proofs that certain aspects of our conclusions are correct. Preferable, however, would be to have found theorems regarding, particularly, the amplitudes of quasi-limit-cycles in the slow-fast model. The book by Berglund and Gentz [20] presents a formal analysis of a very general slow-fast model in the parametric neighbourhood of a Hopf point. They focus on how long solutions to the deterministic slow-fast system remain near what they call the ‘frozen’ solution, which we call the base system solutions, for values of *λ <* 0. They do not, however, consider the magnitudes of either quasi-cycles or quasi-limit-cycles.

A formal analysis of the stochastic slow-fast system that we defined and studied in Section 4 could be an extension of the stochastic analysis of the quasi-cycles, *λ <* 0, in [43] for our class of slow-fast models, and a similar extension of [44] for *λ >* 0 to quasi-limit-cycles for our class of slow-fast models.

We can conclude from our results that it may be theoretically difficult to distinguish, from data, between a noisy fixed point and a noisy limit cycle if either occurs in connection with a Hopf bifurcation. We used a slow-fast system where a slowly varying parameter increased across a Hopf value. Probably equally common in real brains is a decreasing bifurcation parameter value, i.e., a slowly decreasing synaptic efficacy. Although an exploratory example indicated that in our model class there is a smooth transition across the Hopf point in this case, from noisy limit-cycles to quasi-cycles, most theoretical analyses consider only increasing parameter values.

Another theoretical question would be: is there a convenient normal form for our model class? This would involve finding transformations of the various specific model equations into a common form. An example of a normal form for a deterministic model with a Hopf bifurcation is Equation (1). It is possible that a normal form for our model class would resemble this equation but with the addition of a noise term. A common noise term would be equivalent to the one we use in our definition in (2), that is, standard independent Wiener processes. But the transformations of the various specific terms in the different models would all be different and not easy to see immediately, except for the Stuart-Landau model, (8), which is nearly the same as (1).

We need to relate these models in more detail more firmly to real brains, especially regarding expected noise levels, and to explore further their relationship to the various spiking neuron models. This should be fairly easy for models such as the FitzHugh-Nagumo model, which is a simplification of the Hodgkin-Huxley spiking model. This has been done to some extent for the Wilson-Cowan model that, in addition to being derived from ideas about neural spiking, has been shown to summarize a range of properties of a spiking model, including sparse firing and both quasi-cycles and noisy limit cycles [28]. Some progress has been made by Ermentrout [50] in reducing conductance based models with ‘slow synapses’ to firing rate models. His approach is related to the slow-fast systems we study here.

It might seem that another limitation of the present study is that the Gaussian noise we added to our models is perhaps not the most characteristic of brain noise. The power spectral density (psd) of Gaussian (white) noise does not change with frequency, whereas that of brain noise usually decreases with frequency. Indeed, an intriguing aspect of real brains is the fact that they display 1*/f* ^*α*^-like noise spectra at both large (EEG, MEG) and small (single-unit, intracranial recordings) scales, e.g., [51–54]. This type of noise has been said to be related to optimal brain function, and changes with age [55] and with some brain disorders [56]. Quite a few studies have indicated that the psd of the brain noise closely either resembles a Lorentzian with one or more “bumps”, or a bumpy 1*/f* ^*α*^ distribution with 0 *< α ≤* 3, e.g., [51]. Lorentzians are the psds of OU or AR1 processes [57]. In log-log plots Lorentzians are flat at lower frequencies and, after an inflection point that depends on the parameters of the process, descend linearly with frequency, and with a slope of approximately −2, although steeper slopes are possible. Importantly, all of the models studied here have Lorentzian psds with bumps at the inflection point that are associated with the frequency (rotation) of quasi-cycles or noisy (or quasi-) limit cycles. Moreover, as shown in [57], the sum of several Lorentzians with different inflection points is 1*/f* ^*α*^-like for a large range of possible values of *α*. Adding the outputs of several copies of, say, the tanhKang model, with different or changing parameter values, should result in a 1*/f* ^*α*^-like psd, with bumps, that would model the psd of recordings of noisy brain activity. Thus, the models we studied here have the potential to illuminate additional aspects of brain function.

In order to apply our results, further study is needed of how synaptic efficacy changes in real systems and how these changes affect the dynamics of systems. We mentioned earlier a couple of studies of relatively fast-changing synapses [21, 22]. But synapses change over many time scales, from hundreds of milliseconds to minutes or even hours or days, and in both directions, as a function of spike-timing dependence and repeated stimulation. Given how sensitive the dynamics of our model systems are to bifurcation parameters, which can depend on synaptic efficacies as in the tanhKang and Wilson-Cowan examples, and the likelihood of ‘slow’ changes in those efficacies, it is important to understand in detail the ways in which changing synaptic efficacies effect modelled Hopf points in the brain.

